# Model neuron response statistics to natural images

**DOI:** 10.1101/387183

**Authors:** Arvind Iyer, Johannes Burge

## Abstract

To model the responses of neurons in the early visual system, at least three basic components are required: a receptive field, a normalization term, and a specification of encoding noise. Here, we examine how the receptive field, the normalization factor, and the encoding noise impact the model neuron responses to natural images and the signal-to-noise ratio for natural image discrimination. We show that when these components are modeled appropriately, the model neuron responses to natural stimuli are Gaussian distributed, scale-invariant, and very nearly maximize the signal-to-noise ratio for stimulus discrimination. We discuss the statistical models of natural stimuli that can account for these response statistics, and we show how some commonly used modeling practices may distort these results. Finally, we show that normalization can equalize important properties of neural response across different stimulus types. Specifically, narrowband (stimulus- and feature-specific) normalization causes model neurons to yield Gaussian-distributed responses to natural stimuli, 1/f noise stimuli, and white noise stimuli. The current work makes recommendations for best practices and it lays a foundation, grounded in the response statistics to natural stimuli, upon which principled models of more complex visual tasks can be built.

## Introduction

As interest intensifies in understanding natural signals in vision and neuroscience, it becomes increasingly important to develop a clear picture of how neural systems and their constituent components respond to real-world (photographic) images. Characterizing the statistical properties of these responses is vital for building principled models of visual processing, especially given the increasing reliance of vision and visual neuroscience on probability theory (Knill & Richards, 1996). Over the past two decades, there have been many neurophysiological (Baudot et al., 2013; Burkhardt, Fahey, & Sikora, 2006; Butts et al., 2010; Felsen, Touryan, Han, & Dan, 2005; Lesica et al., 2007; Talebi & Baker, 2012; Weliky, Fiser, Hunt, & Wagner, 2003) and computational (Brady & Field, 2000; Burge & Geisler, 2014; 2015; Clatworthy, Chirimuuta, Lauritzen, & Tolhurst, 2003; Lyu & Simoncelli, 2008; 2009b; Sebastian, Abrams, & Geisler, 2017; Tadmor & Tolhurst, 2000; Wainwright & Simoncelli, 2000) attempts to address this issue.

We report a large-scale analysis of model neuron response statistics to natural images. To model neural responses, at least three basic components are required: a receptive field, a normalization term, and a specification of encoding noise. The receptive field specifies the neuron’s preferred stimulus feature, and indicates how inputs are weighted and summed (i.e. pooled) across space to determine the response (Hubel & Wiesel, 1962; 1968). The normalization term specifies how gain control is implemented to prevent the neuron’s response from exceeding its dynamic range (Albrecht & Geisler, 1991; Heeger, 1992). The encoding noise specifies the uncertainty in the response to repeated presentations of the same stimulus (Tolhurst, Movshon, & Dean, 1983a).

We examine how methods for modeling the receptive field, the normalization factor, and the encoding noise impact the response statistics to natural images and the signal-to-noise ratio for stimulus discrimination. We focus our analysis on model neurons with oriented receptive fields like those in early visual cortex. We show that feature-specific normalization of each stimulus, with an easy to compute normalization factor, yields model neuron responses to natural images that are Gaussian distributed. We show that the stimulus-driven response variance is invariant to the scale of the preferred feature. We also show that responses with these statistics nearly maximize the signal-to-noise ratio for stimulus discrimination given the noisy neural response. Furthermore, we show that subtle variants of the standard response model, which are used across the vision and computational neuroscience communities, have an important impact on the results described above. To achieve scientific consensus on basic facts about natural stimulus processing early in the visual system, it is important to understand how variants of the standard response model impact response statistics.

## Results

How do common receptive field modeling choices impact response statistics and natural image discrimination? To investigate, we first establish the relation between response variability and stimulus discriminability. Second, we describe a common model of neural response in early visual cortex and discuss how two different forms of normalization impact the response statistics and impact natural stimulus discriminability. Third, we discuss the statistical models of natural stimuli that can account for these results.

### Stimulus discriminability from neural response

Consider a model neuron that produces a particular response distribution across tens of thousands of natural stimuli. Any early visual representation must be capable of distinguishing two arbitrary stimuli from one another. How well can a neuron distinguish to arbitrary stimuli from the ensemble of natural stimuli? To assess the signal-to-noise ratio (SNR) for stimulus discrimination given the responses of a particular neuron, we compute the discriminability of two arbitrary stimuli, randomly sampled from the natural distribution of stimuli.

The discriminability (i.e. d-prime) of any two stimuli based this neuron’s response is given by

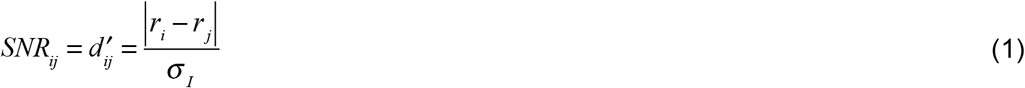

where *r_i_* and *r_j_* represent the expected model neuron responses to two randomly sampled stimuli and *σ_1_* represents internal encoding noise.

The response distribution *p*(*r*) to natural stimuli has a critical impact on discriminability. Under the assumption that responses to natural stimuli are Gaussian distributed 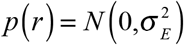 the expected discriminability across all stimuli is given by

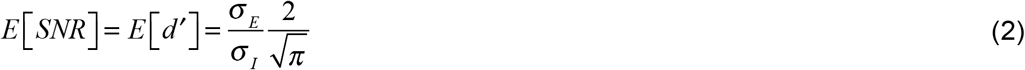

where *σ_E_* is the stimulus-driven (external) response variation (e.g. external noise; see Fig. 1) and the expectation is taken across all stimulus pairs (see Supplement). (If responses are Laplace distributed, the expected discriminability is given by 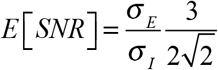; see Supplement). For an arbitrary response distribution, expected SNR can be computed using numerical methods. The fact that the stimulus-driven standard deviation is in the numerator of Eq. 2 indicates that greater stimulus-driven response variation yields better stimulus discriminability.

**Figure 1.**
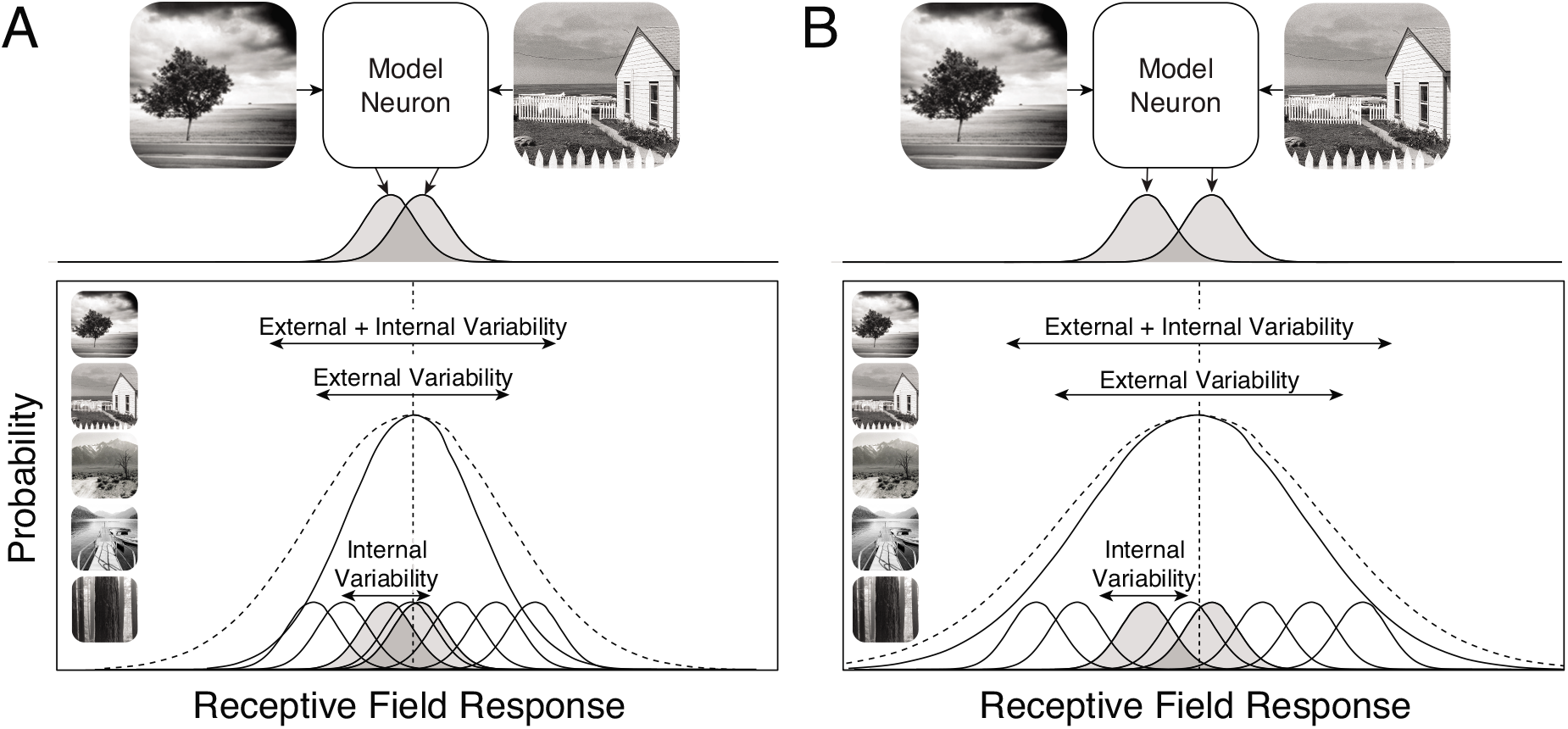
Stimulus-driven response variability and the signal-to-noise ratio for stimulus discrimination. **A** Model neuron yielding low stimulus-driven response variance. Two natural images elicit encoding-noise-corrupted responses (shaded bell curves) that are hard to discriminate based on the responses (top). Low stimulus-driven response variability to the natural image ensemble is associated with poor signal-to-noise for stimulus discriminability (bottom). Two randomly stimuli will be hard to discriminate on average. **B** Model neuron with high stimulus-driven response variance. The same two random stimuli are now easier to discriminate, as will randomly sampled images from the natural image ensemble.

Increased variance is usually associated with poorer stimulus discriminability (Ernst & Banks, 2002), so Eq. 2 deserves further explanation. The source of the response variance is critical for determining whether it helps or hurts discrimination. When response variance is primarily due to noise, stimulus discriminability is poor. When response variance is primarily stimulus-driven, stimulus discriminability is high. For example, if the sensory afferents to a model neuron’s receptive field are severed, the stimulus-driven response variance will be zero, and discriminating stimuli based on its response would be impossible. Fig. 1A shows a model neuron with low stimulus-driven response variance. Fig. 1B shows a model neuron with high stimulus-driven response variance. The top row shows how encoding noise (shaded bell curves) limits discriminability for two natural stimuli. The bottom row shows the impact of stimulus-driven variability across the image ensemble. High stimulus-driven variability will tend to yield larger differences between the expected responses to two random stimuli. Thus, discriminability improves when the source of the response variability is external and stimulus-driven, and discriminability deteriorates when the source of the response variability is internal and due to noise. It should be noted that the design of the visual system is surely driven by tasks more sophisticated than stimulus discrimination but for present purposes, it is a useful task around which to organize our discussion.

### Model neuron responses

Responses of neurons in early visual cortex are commonly modeled as arising from a series of processing stages that are constrained by known properties of neural response in the early system (Fig. 2A). First, a linear receptive field filters the stimulus to yield the linear response. Next, the linear response is normalized by a factor that is determined by local properties of the stimulus or by the responses of other neurons in a local pool (Albrecht & Geisler, 1991; Heeger, 1992); the normalized response is called the ‘response drive’. Last, the response drive is corrupted by encoding noise. Many models of neural response also incorporate response rectification and a static output non-linearity. We consider the effect of these non-linearities in the discussion, but our primary analysis is focused on the response drive statistics.

**Figure 2.**
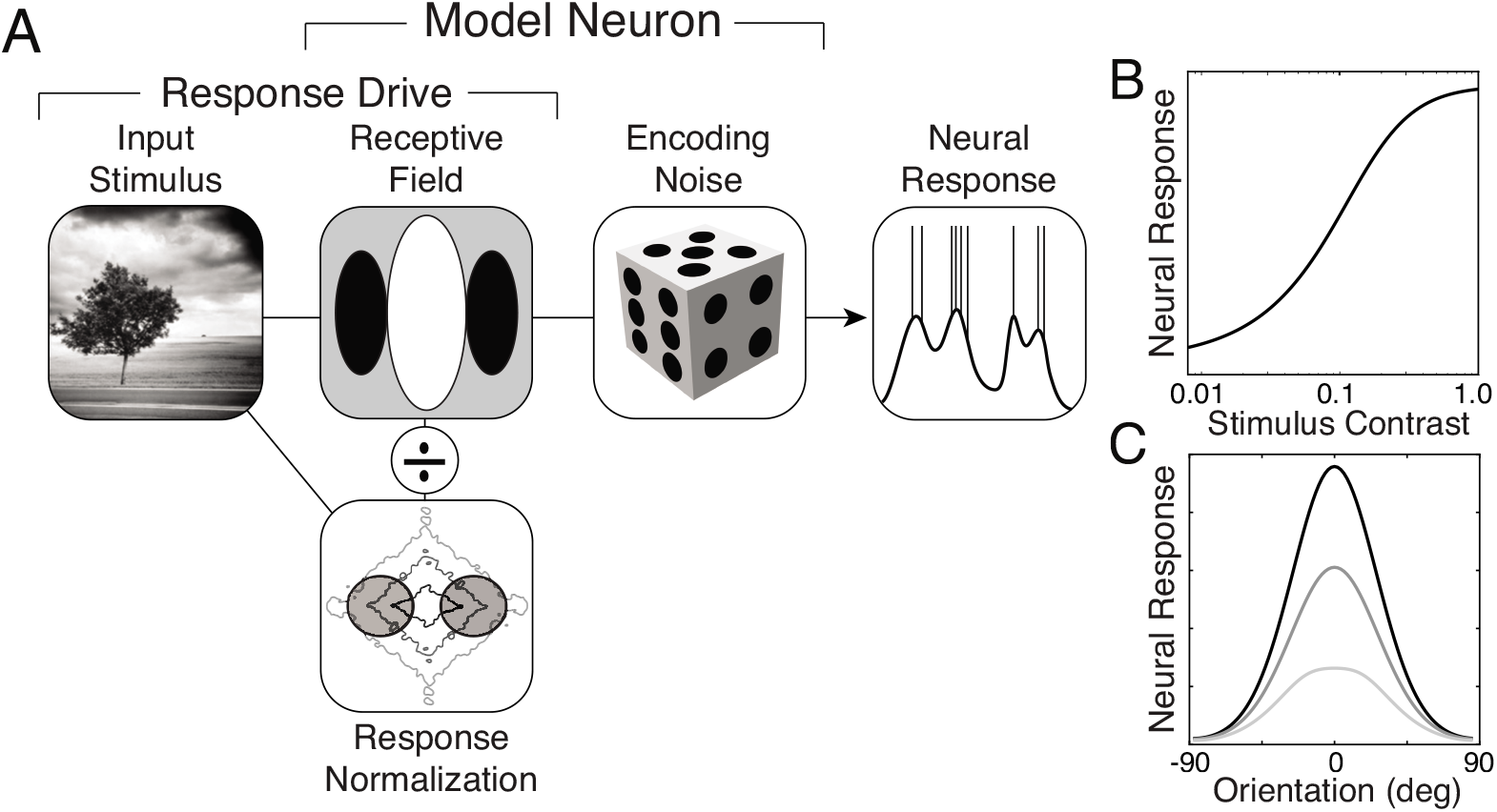
Model neuron. **A** Critical stages of neural response model: linear filtering, response normalization, and encoding noise. The stimulus is encoded by a linear filter, normalized by a portion of the stimulus energy, and then corrupted by encoding noise. The model neuron response provides a prediction of intracellular voltage or spike rate, depending on circumstances. **B** Contrast response function for the receptive field’s preferred stimulus: a vertical Gabor. **C** Orientation tuning function for three different spatial frequencies: stimulus frequency equals the preferred spatial frequency (black), 2x the preferred spatial frequency (dark gray), and 3x the preferred spatial frequency (light gray).

More specifically, the response *R* to a particular stimulus is given by

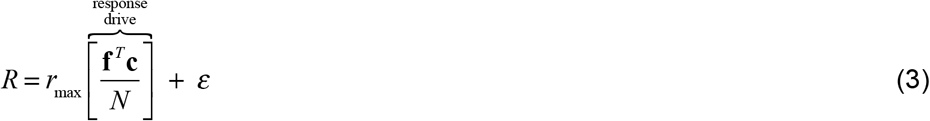

where *r*_max_ is the neuron’s maximum response, **f** is the receptive field, **c** is a contrast stimulus (possibly corrupted by input noise), *N* is the normalization factor, and 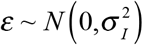 is encoding noise. The standard deviation *σ*_1_ of the internal encoding noise can be constant or it can scale with the mean response (i.e. ‘Poisson-like’). The receptive field is assumed to have a vector magnitude (i.e. L2 norm) of 1.0.

The maximum response is set to a constant for all model neurons. In individual simple cells, maximum firing rate is thought to be independent of preferred spatial frequency, orientation, and other stimulus preferences. It has been observed that overall firing rate in cortex tends to decrease as spatial frequency increases. But this decrease in overall firing rate is likely to be a population effect due to a non-uniform distribution of spatial frequency preferences in cortex, to sampling bias for low spatial frequencies in neuroscience studies, or both (De Valois, Albrecht, & Thorell, 1982a; Foster, Gaska, Nagler, & Pollen, 1985; Victor, Purpura, Katz, & Mao, 1994). Thus, in the current paper, and without loss of generality, for all receptive fields we assume r_max_ equals 1.0. We focus on characterizing the statistics of the response drive **f**^*T*^**c**/*N* to natural stimuli.

The response model in equation 3 enforces the limited dynamic range of neural response in cortex and helps describe the shape of the contrast response functions of neurons in cortex (Fig. 2B). The response model also accounts for the invariance of the shapes of orientation tuning curves to Gabor or grating stimuli having different spatial frequencies (Fig. 2C).

### Receptive field

A receptive field is a function that weights and sums inputs across space and time to determine a neuron’s response. Responses increase when receptive field locations having positive weights are stimulated with input increments and decrease when stimulated with input decrements. The opposite happens with locations having negative weights. In early visual cortex, simple cell receptive fields are often modeled as having the shape of a Gabor—a cosine wave windowed by a Gaussian envelope (Jones & Palmer, 1987a; 1987b). Gabor receptive fields are orientation and spatial frequency selective; the selectivity is commonly quantified by the bandwidth. The orientation bandwidth specifies the range of input orientations that can elicit a response. The median orientation bandwidth in cortex is 42° (De Valois, Yund, & Hepler, 1982b). The spatial frequency bandwidth specifies the range of input spatial frequencies that can elicit a response. The distribution of simple cell bandwidths in cortex ranges between 0.8 to 1.8 octaves at half-height, with a median bandwidth of 1.2 octaves (De Valois, Albrecht, & Thorell, 1982a). We will characterize the response statistics of model neurons with vertically oriented Gabor receptive fields having the median orientation bandwidth of 42° and spatial frequency bandwidths that span the same range as simple cell receptive fields in cortex (octave bandwidths = 0.8, 1.2, and 1.8; Fig. 3AB). We examine model neuron responses having receptive fields with preferred spatial frequencies between 2 and 8 cpd.

**Figure 3.**
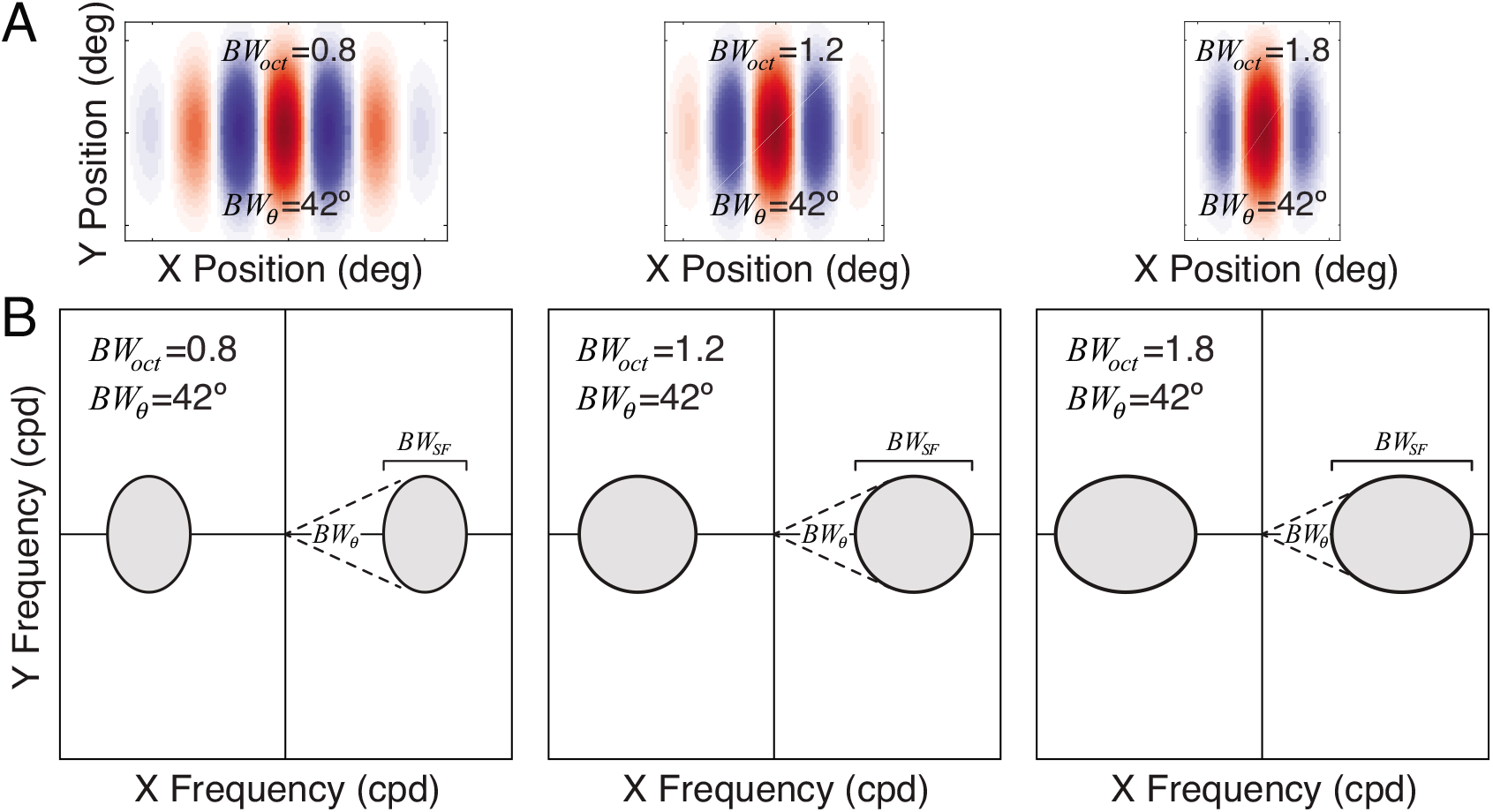
Gabor receptive fields and amplitude spectra. **A** Gabor receptive fields with octave bandwidths of 0.8, 1.2, and 1.8, and orientation bandwidths of 42°. Different octave bandwidths correspond to preferred features with different aspect ratios (see Methods). **B** Amplitude spectra of Gabor receptive fields. Orientation bandwidth *BW_θ_* is the polar angle spanned by the amplitude spectrum at half-height. Spatial frequency bandwidth *BW_SF_* = *f_hi_* – *f_lo_* is the range of frequencies spanned by the spectrum, where *f_hi_* and *f_lo_* are the high and low frequencies at half-height. Octave bandwidth *BW_oct_* = log_2_ (*f_hi_*/*f_lo_*) is the log-base-two ratio of the frequencies.

Early models of neural response proposed that response drive is a linear function of the input stimulus (Campbell, Cleland, Cooper, & Enroth-Cugell, 1968; Hubel & Wiesel, 1962; 1968). The linear receptive field responses *R_lin_* = **f**^*T*^**c** to natural stimuli are nicely approximated by a generalized Gaussian with tails heavier than a Laplace distribution (Fig. S1). Previous analyses of linear responses have reported similar findings (Wainwright & Simoncelli, 2000). We do not focus on the linear responses because real neurons include response normalization.

### Response normalization

The linear model explains neural responses in some regimes, but it is insufficiently rich to capture neural response properties over a wide range of stvimulus conditions. Response normalization was originally proposed to account for the limited dynamic range of neurons in early visual cortex (Albrecht & Geisler, 1991; Heeger, 1992). Evidence for response normalization has been observed in primate retina, lateral geniculate nucleus, and early visual cortex (Albrecht & Geisler, 1991; Benardete, Kaplan, & Knight, 1992; Carandini, Heeger, & Movshon, 1997; Chander & Chichilnisky, 2001; Heeger, 1992; Mante, Bonin, & Carandini, 2008; Mante, Frazor, Bonin, Geisler, & Carandini, 2005; Nishimoto, Ishida, & Ohzawa, 2006; Shapley & Victor, 1978; Solomon, Peirce, Dhruv, & Lennie, 2004). In more recent years, normalization has been proposed to occur in higher cortical areas and be associated with computations underlying diverse behavioral phenomena (Carandini & Heeger, 2012). We examine how two types of response normalization—broadband normalization and narrowband normalization—impact the responses of model neurons to natural stimuli.

Broadband normalization is stimulus-specific but feature-independent. With broadband normalization, the model neuron responses are normalized by all the stimulus contrast in a local image region at the receptive field location, regardless of its preferred feature (Carandini et al., 1997); all orientations and spatial frequencies normalize the linear response. The broadband normalization factor is

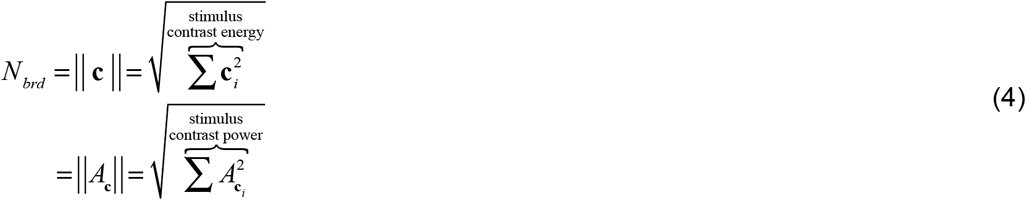

where **c** is a (possibly noisy) Weber contrast stimulus, *A_c_* is the amplitude spectrum of the contrast stimulus, and the L2 norm operator ‖·‖ gives the square root of the sum of squares. Parseval’s theorem guarantees that the total energy of the contrast stimulus equals the total power of its amplitude spectrum (Fig. 4A). Note that if the contrast stimulus is noisy (e.g. corrupted by pixel noise), the broadband normalization factor equals 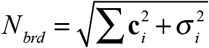 to very close approximation, where *σ* is the standard deviation of the input noise (Burge & Geisler, 2014).

**Figure 4.**
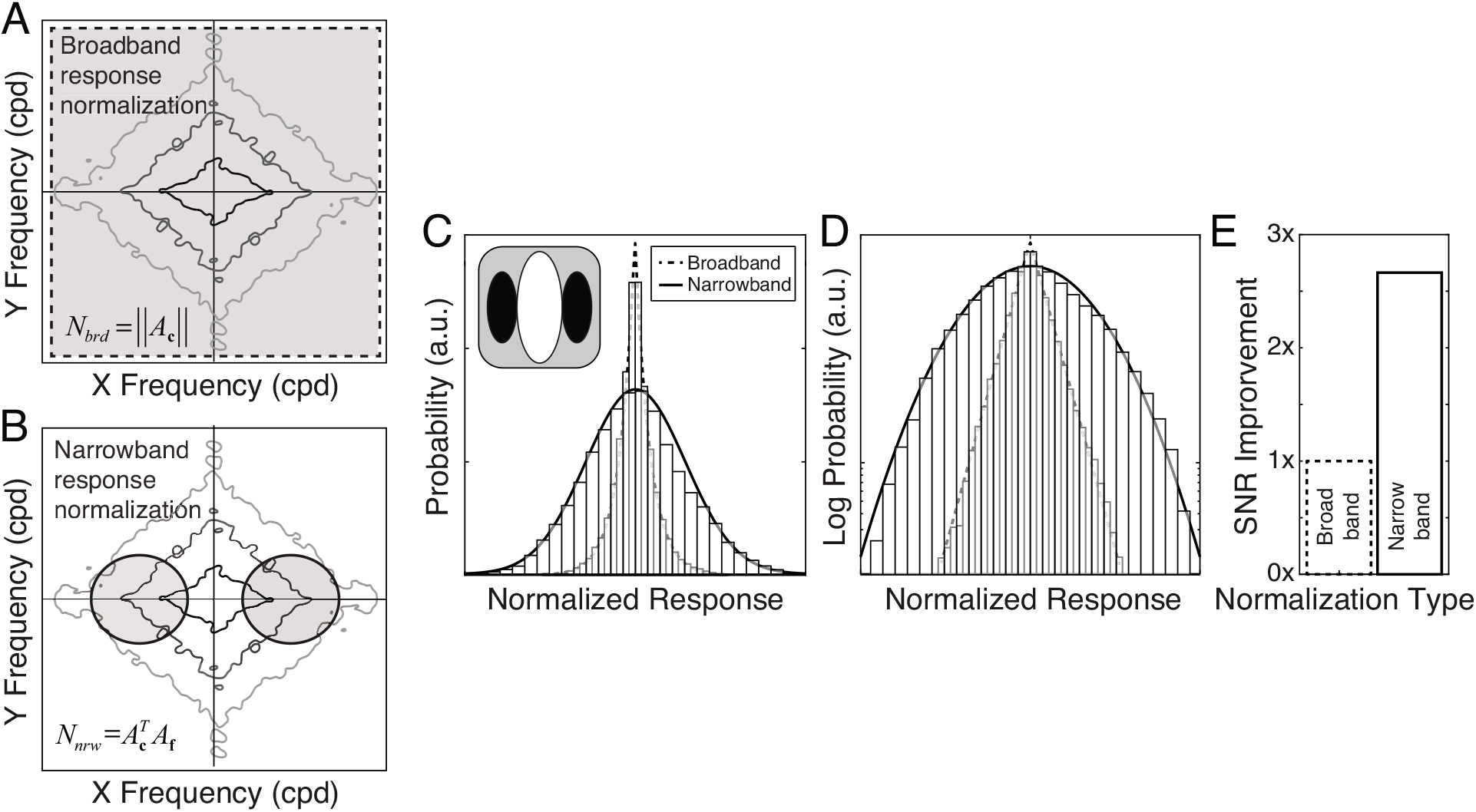
Broadband vs. narrowband normalization with a Gabor-shaped receptive field. **A** Broadband normalization uses all the stimulus contrast (gray area) to normalize the linear receptive field response, regardless of orientation and spatial frequency. The diamond shaped contours represent the amplitude spectrum of an individual natural image patch. **B** Narrowband normalization uses only the stimulus contrast in the pass band of the receptive field (gray area) to normalize the linear receptive field response. **C** Probability of model neuron responses. With broadband normalization, receptive field responses to natural stimuli are highly non-Gaussian, and are nicely approximated by a Laplace distribution (dashed curve). With narrowband normalization, the same receptive field yields responses to natural stimuli that are well described by a Gaussian (solid curve). **D** Same responses as in C, but with the y-axis showing log-probability over three orders of magnitude. **E** Factor improvement in signal-to-noise (SNR) for stimulus discriminability with narrowband vs. broadband normalization. Results are shown for a vertically oriented Gabor with an orientation bandwidth of 42° and an octave bandwidth of 1.2. Similar results are obtained for other receptive fields.

Narrowband normalization is stimulus-specific and feature-dependent. With narrowband normalization, the model neuron responses are normalized by the stimulus contrast in the pass band of the receptive field, which means that the stimulus features that contribute most prominently to the normalization factor are those that approximately match the preferred feature (Fig. 4B). For example, if the receptive field’s preferred stimulus is a vertically oriented Gabor with a carrier frequency of 4 cpd, the responses are normalized primarily by features that are near vertical and near to 4 cpd. The narrowband normalization factor is given by

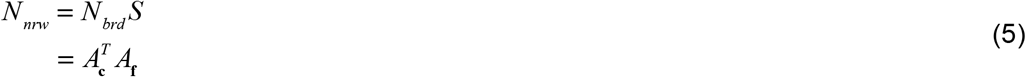

where 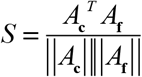 is the phase-invariant similarity, the cosine similarity between the stimulus and receptive field amplitude spectra (Sebastian et al., 2017). (The amplitude spectrum of the receptive field *A*_**f**_ is assumed to have an L2 norm of 1.0.) Similarity is thus constrained to take a value between 0 and 1, which means that the narrowband normalization factor is always less than or equal to the broadband factor.

Broadband-normalized responses *R_brd_* ∞ **f**^*T*^**c**/*N_brd_* to natural stimuli are highly non-Gaussian. The Laplace distribution provides an excellent fit to the broadband responses for all preferred spatial frequencies and octave bandwidths (Fig. 4C,D; Fig. S2). Narrowband-normalized responses *R_nrw_* ∞ **f**^*T*^**c**/*N_nrw_* differ from broadband responses in two important ways. First, natural-stimulus-driven response standard deviation *σ_E_* is approximately two and a half times higher for narrowband than broadband responses. Second, narrowband normalization yields response distributions that are very nearly Gaussian (Fig. 4C,D; Fig. S3). Related findings have been reported by other groups (Burge & Geisler, 2014; 2015; Jaini & Burge, 2017; Lyu & Simoncelli, 2008; 2009a; Sebastian et al., 2017; Wainwright & Simoncelli, 2000).

Relative to broadband normalization, narrowband normalization also improves signal-to-noise for stimulus discrimination by nearly three times, assuming constant encoding noise (Fig. 4E). The improvement in signal-to-noise is mediated both by the more Gaussian (lighter-tailed) response distributions (Fig. S4A), and by the increased stimulus-driven response variability (Eq. 2). Poisson-like encoding noise, which is more like response noise in cortex (Tolhurst, Movshon, & Dean, 1983b), yields very similar results (Fig. S4B-D).

Why does narrowband normalization increase stimulus-driven response variance relative to broadband normalization? Because the narrowband normalization factor is always less than or equal to the broadband normalization factor (Fig. 5A; Eqs. 4,5). Therefore, across many stimuli, the response distribution will tend to have larger variance with narrowband normalization. Why does narrowband normalization result in more Gaussian responses than broadband normalization? Because narrowband normalization causes small broadband responses to be amplified significantly, and leaves large broadband responses relatively unchanged (Fig. 5B). For example, if the stimulus is a poor match to the receptive field (i.e. the broadband response approaches 0.0), it is likely that only a small proportion of the stimulus contrast is in the pass band of the receptive field. This, in turn, means that the narrowband normalization factor will be quite small compared to the broadband factor, which will increase the proportion by which the narrowband response is amplified relative to the broadband response (Eq. 5; Fig. 5B,C). On the other extreme, if the stimulus is a *perfect* match to the receptive field, the broadband response equals r_max_, and all the stimulus contrast must be in the pass band of the receptive field. The narrowband and broadband normalization factors will thus be identical, and the narrowband response will equal the broadband response. Intermediate broadband responses are amplified by an intermediate amount (Fig. 5C). These effects mediate the differences in the shapes of the broadband and narrowband response distributions (Fig. 5D).

**Figure 5.**
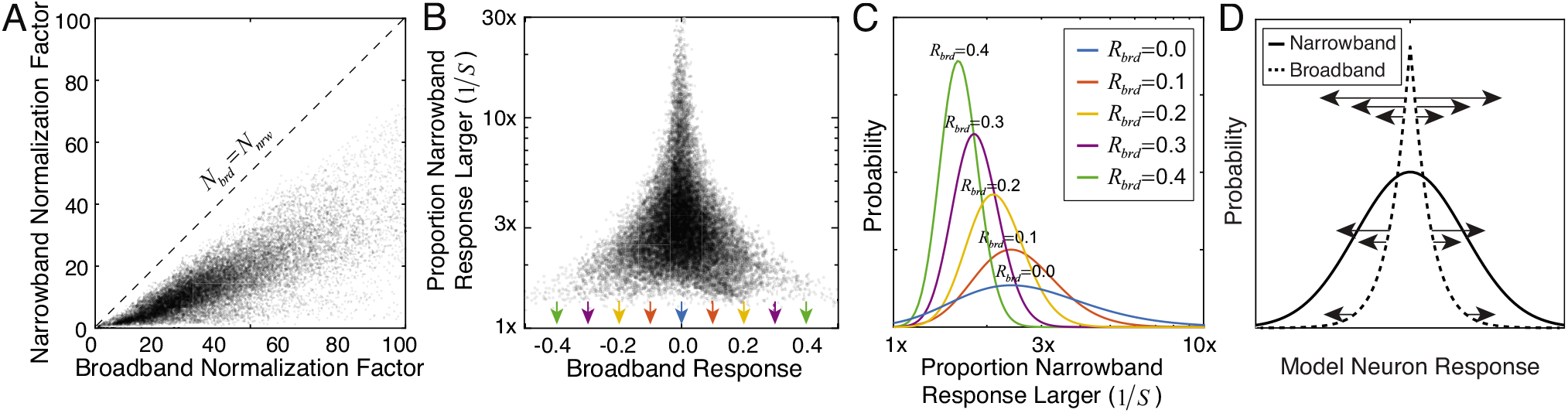
Broadband vs. narrowband normalization. **A** The narrowband normalization factor is smaller than the broadband normalization factor for each stimulus. **B** Proportion that the narrowband response is larger than the broadband response *R_nrw_*/*R_brd_* as a function of the broadband response. Large proportions occur only for small broadband responses, accounting for why narrowband normalization increases the Gaussianity of the model neuron responses. **C** Distribution of the proportional increase in the narrowband response relative to the broadband response, conditioned on different absolute values of the broadband response (colors), as fit by inverse gamma distributions (Fig. S5). The proportion is equivalent to inverse similarity. Arrows in B mark the absolute values of broadband response upon which the proportions are conditioned. **D** Schematic showing why the relationship between the narrowband and the broadband responses contributes to the increased Gaussianity of the model neuron responses. The data in A-C are for a vertically oriented cosine-phase Gabor receptive field with an orientation bandwidth of 42°, an octave bandwidth of 1.2, and a preferred frequency of 2 cpd. Results are similar for all receptive fields.

The proportional increase *R_nrw_*/*R_brd_* of the narrowband response vs. the broadband response depends strongly on the broadband response (Fig. 5B). Figure 5C shows conditional distributions of proportional increase for five different absolute values of the broadband responses, as fit by inverse gamma distributions (Fig. S5). When stimuli are narrowband normalized, small broadband responses are amplified more than large broadband responses.

There is an additional point worth making. The results presented in Fig. 4C,D suggest that the broadband-normalized responses can be represented as a Gaussian scale mixture of random variables. When considering natural images, the input contrast image c is a random variable. It follows that the broadband-normalized response *R_brd_*, the narrowband-normalized response *R_nrw_*, and the phase-invariant similarity *S* are all random variables. By combining equations 4–5, it is easy to show that these variables have the following relationships

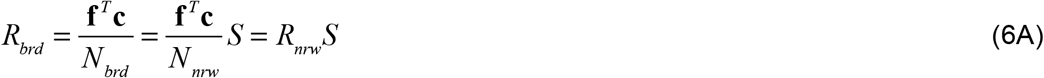

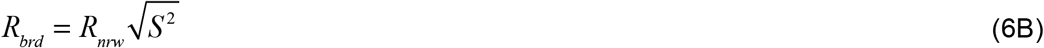

Equation 6B implies th at the broadband-normalized responses are distributed as *R_brd_* ~ *N*(o,S^2^) because the narrowband-normalized responses are approximately zero-mean Gaussian (see Fig. 4C,D). Furthermore, given that the broadband responses are approximately Laplace-distributed (see Fig. 4C,D), equation 6B also implies that the square of the phase-invariant similarity should be approximately Gamma distributed. This is because the Laplace distribution can be represented as a Gaussian scale mixture when the mixing distribution (i.e. the variance of the Gaussian) is Gamma distributed with a shape parameter of 1.0 (i.e. an Exponential distribution *S* ^2^ ~ Γ (*α* = 1,*β*) = *Exp* (*β*)) (Ding & Blitzstein, 2018). Figure 6B,C shows that *S*^2^ is indeed nicely approximated by a Gamma distribution with a shape parameter of 1.4, which is Exponential to close approximation. Thus, normalizing the broadband responses by the similarity yields Gaussian-distributed narrowband responses.

**Figure 6.**
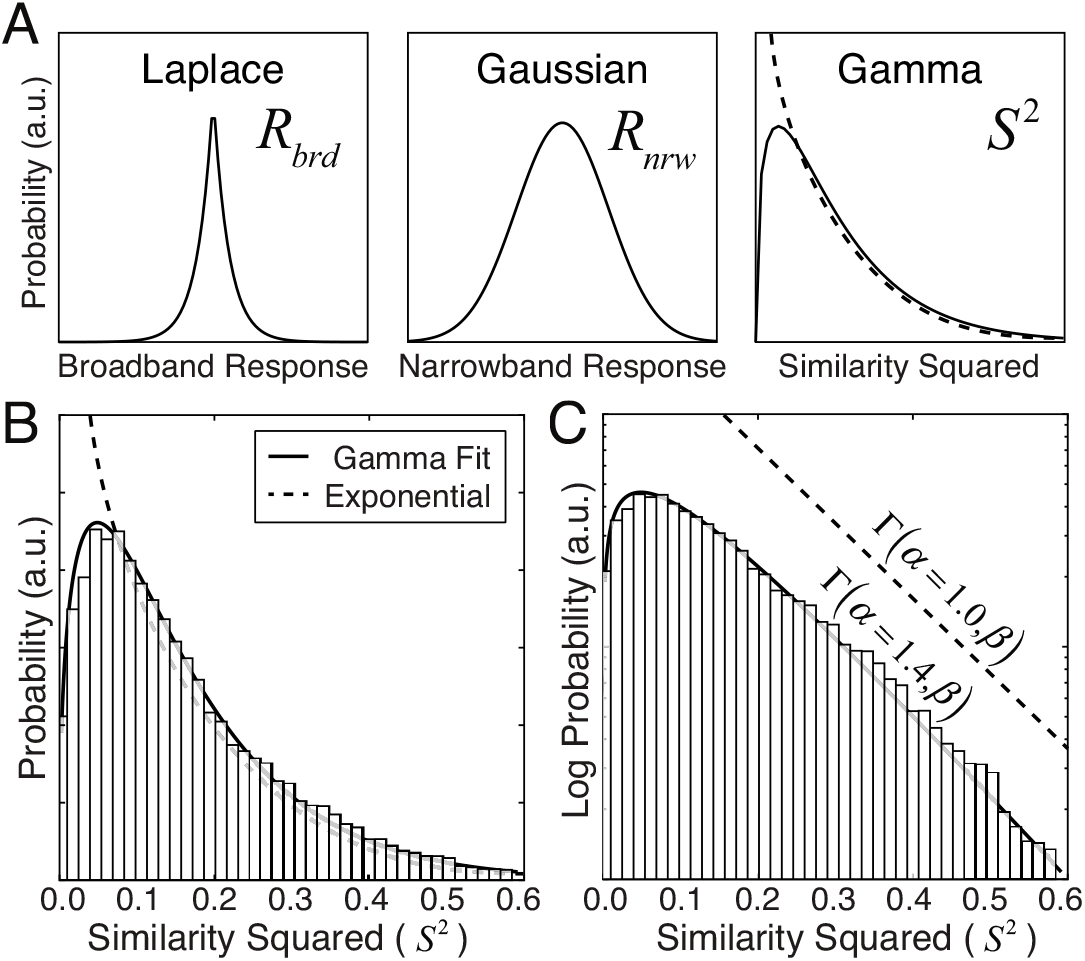
Broadband responses represented as a Gaussian scale mixture. **A** Laplace-distributed broadband responses can be expressed as a scale mixture of Gaussian narrowband responses with a Gamma (i.e. Exponential) distributed mixing variable. **B** Squared similarity across all natural stimulus (bars) and a Gamma distribution fit via maximum likelihood (solid curve); an Exponential distribution (a gamma distribution with a shape parameter of 1.0) is shown for reference (dashed curve). **C** Same data as A on a log-probability axis spanning two orders of magnitude with Gamma and Exponential fits. The exponential distribution is shifted vertically to reduce clutter. These results data are for a vertically oriented cosine-phase Gabor receptive field with an orientation bandwidth of 42°, an octave bandwidth of 1.2, and a preferred frequency of 2 cpd. Results are similar for all receptive fields.

Lyu & Simoncelli (2008) also modeled linear filter responses to natural images as a Gaussian scale mixture. Specifically, they modeled the linear (un-normalized) filter responses as a Gaussian scale mixture. They estimated the value of the mixing random variable, from the joint responses of a large bank of multi-scale filters. One potential advantage of the work presented here is that the narrowband normalization factor (i.e. the value of the mixing random variable) can be computed directly from the amplitude spectra of the stimulus and the receptive field (Eq. 5). Being able to compute the normalization factor directly from the stimulus may make stimulus- and feature-specific normalization easier to implement for some computational investigations.

### Normalization pooling region

The receptive field specifies how inputs are weighted and pooled across space to determine the stimulus drive (i.e. **f**^*T*^**c**) to neural response. Receptive fields are typically modeled by a matrix of positive and negative weights that determine how the value of each stimulus pixel contributes to the response (see Fig. 3). Here, we ask how the visual angle spanned by the receptive field weight matrix impacts the response statistics of model neurons to natural stimuli. The visual angle spanned by the receptive field weight matrix impacts the statistics because it determines the stimulus region from which the normalization factor is computed (Eqs. 4, 5; see below).

Consider two sets of neurons employing narrowband contrast normalization having receptive fields with Gabor-shaped preferred features. In the first set, the visual angle spanned by the receptive field weight matrix becomes increasingly mismatched to its preferred feature with increases in preferred spatial frequency (Fig. 7A). In the second set, the visual angle spanned by the receptive field weight matrix is matched to the preferred feature regardless of its spatial frequency (Fig. 7B). These two sets of model neurons produce very different sets of response distributions to natural stimuli. When the weight matrix and preferred feature are matched (i.e. span the same visual angle), the normalization factor is computed from the same image region that drives the linear receptive field response, and the response distributions have constant variance and are approximately Gaussian for all preferred spatial frequencies. When the weight matrix and preferred feature are mismatched, the normalization factor is computed from an image region larger than the preferred feature, the response variance decreases with the inverse frequency (1/f) of the preferred feature and the response distributions become less Gaussian (Fig. 7C). These results are summarized by the stimulus-driven response standard deviation and kurtosis (Fig. 7D,E). Note also that for the largest mismatch considered here, the narrowband response distribution is well approximated by a Laplace distribution. Thus, for large mismatches the benefits of narrowband response normalization are surrendered. (See Supplement for results with broadband normalization; Fig. S6.)

**Figure 7.**
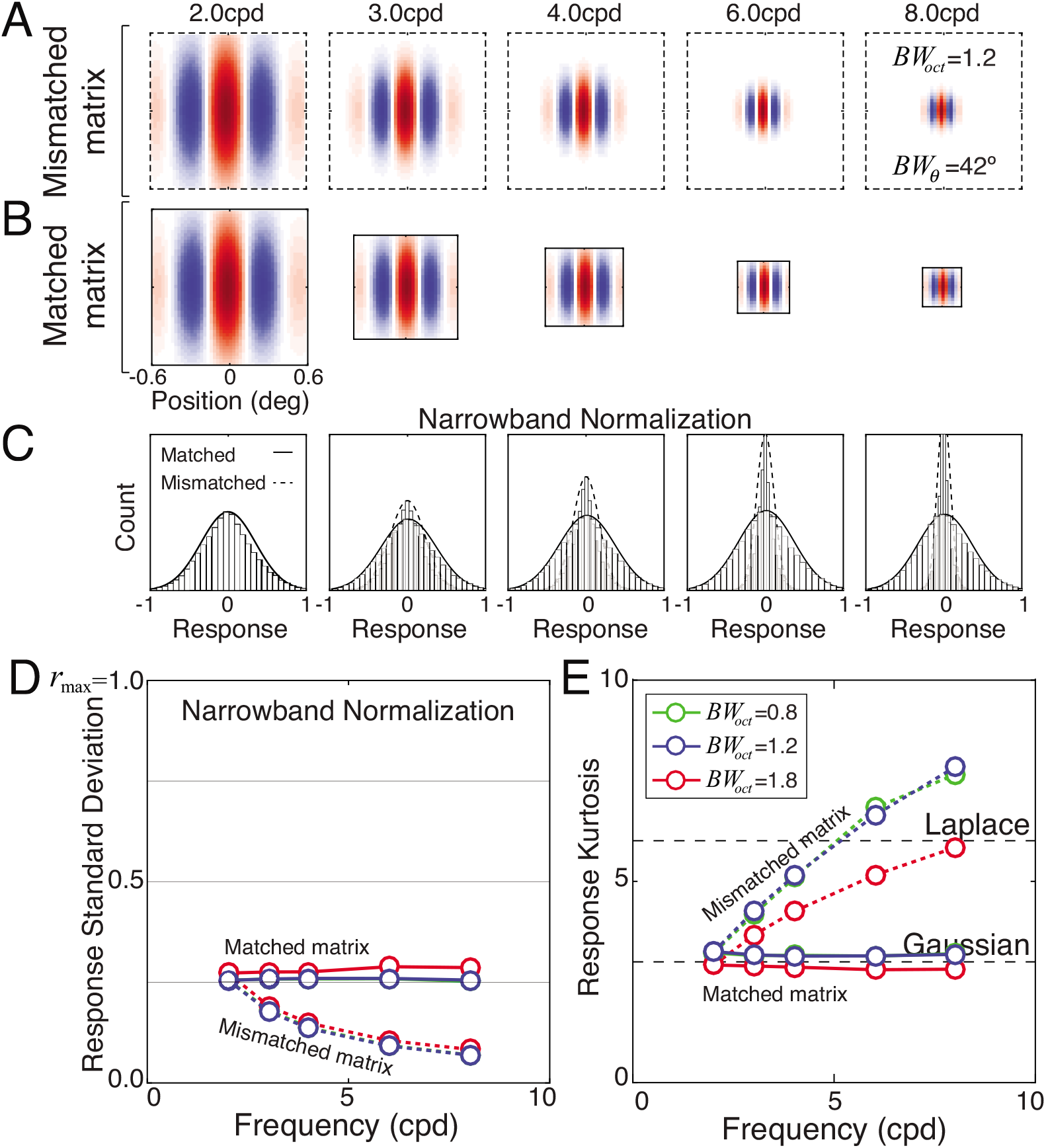
Narrowband response statistics with receptive field weight matrices that are **A** mismatched and **B** matched to the preferred feature: a vertically oriented Gabor with 1.2 octave bandwidth and 42° orientation bandwidth. **C** Response distributions from matched and mismatched matrices. Matched weight matrices (solid curves) yield response distributions that are invariant to the scale of the preferred feature. Mismatched weight matrices (dashed curves) yield response distributions that change shape and variance with the magnitude of the mismatch. **D** Response standard deviation as a function of preferred spatial frequency for octave bandwidth (colors). Stimulus-driven response variance is constant with preferred frequency when the matrix is matched to the preferred feature. When the matrix is mismatched, response variance decreases with the magnitude of the mismatch. **E** Response kurtosis is the same as a Gaussian with matched weight matrices, but increases with the amount of mismatch.

The primary finding, however, is that with narrowband normalization and matched receptive fields, responses are approximately Gaussian, mean-zero, and strikingly invariant to the scale of the preferred feature. Regardless of the preferred spatial frequency, the response is equally statistically reliable. Furthermore, the results indicate that it is exceedingly rare for a neuron to be stimulated by its preferred feature in natural scenes. The stimulus-driven response standard deviation equals approximately 25% of the maximum response, which means that less than one natural stimulus in 10,000 will cause a response within 5% of the neuron’s maximum response.

How should these results inform our thinking about neurophysiological processing of real-world signals? The majority of single-unit neurophysiology has focused on characterizing the stimuli to which individual neurons respond most strongly. We owe much of our knowledge about the response properties of V1 neurons to this approach. But the real-world stimuli that drive neural responses most strongly occur only very rarely. To understand the coding problems faced by nervous systems in natural viewing, it is critical to understand how neurons respond to the stimulus ensemble encountered in the real world.

**Figure 8.**
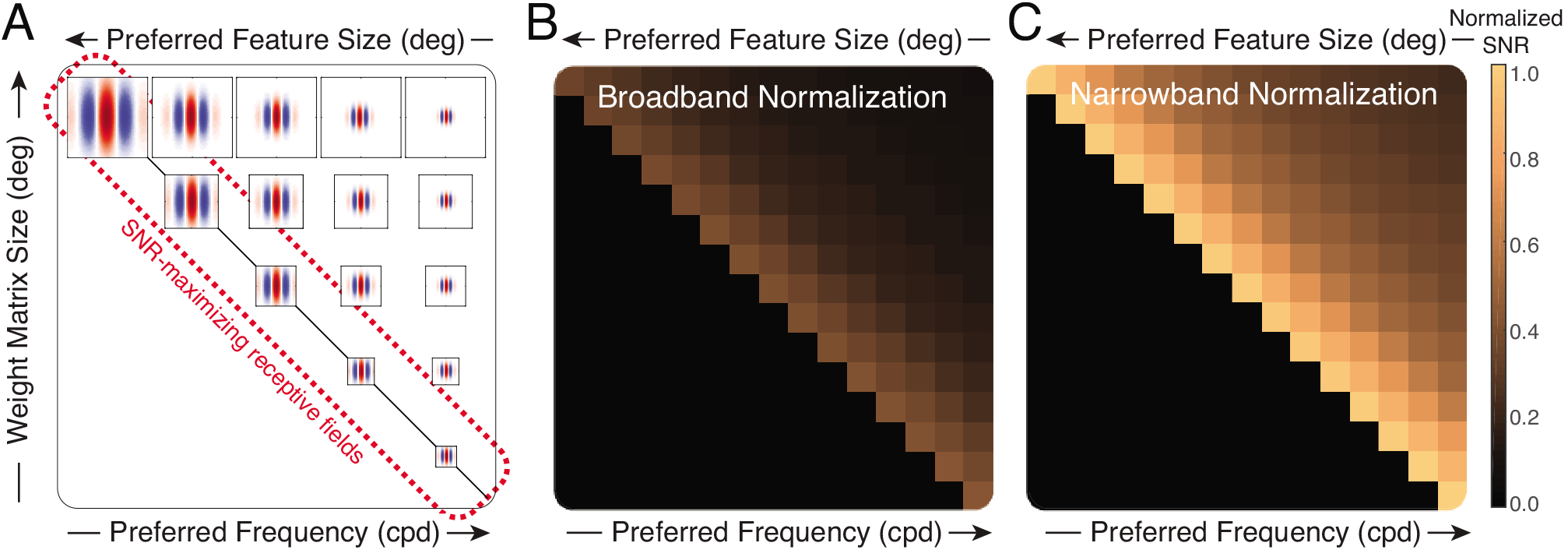
The impact of different receptive field modeling choices on signal-to-noise ratio (SNR). **A** Model receptive fields maximize signal-to-noise ratio (SNR) for stimulus discrimination when the visual angle subtended by the receptive field weight matrix matches the visual angle of the preferred feature (on-diagonal receptive fields). When the visual angles are mismatched, SNR is lower. **B** Normalized SNR with broadband normalization. **C** Normalized SNR with narrowband normalization as a function of preferred spatial frequency and the visual angle spanned by the weight matrix.

To summarize the impact of matching the weight matrix to the preferred feature, we plot the signal-to-noise ratio for stimulus discrimination for a range of preferred features and weight matrix sizes (Fig. 8A). There is a substantial advantage i) for narrowband over broadband normalization, and ii) for matching the visual angle of the receptive field weight matrix to the visual angle of the preferred stimulus feature (Fig. 8B,C). Thus, to maximize the signal-to-noise ratio for stimulus discriminability and to achieve scale invariant response statistics to natural stimuli, one should perform narrowband normalization with weight matrices that match the receptive field’s preferred feature.

As mentioned earlier, these effects occur because of the nonlinear effects of response normalization. The visual angle spanned by the weight matrix determines the size of the image region from which the normalization factor is computed. With mismatched matrices, the normalization factor is determined from the stimulus contrast in an image region larger than the preferred feature, that will likely contain spatial frequencies lower than the preferred frequency. Natural images have 1/f amplitude spectra (D. J. Field, 1987); contrast energy at frequencies lower than the preferred frequency is likely to dominate and substantially increase the value of the normalization factor, thereby decreasing the normalized response. As the mismatch increases, the decrease in the normalized response becomes more pronounced, reducing the stimulus-driven response variability associated with high frequency features. The normalization factor should therefore be determined from the same image region that is selected for by the preferred feature.

Another way to understand the effects is to consider two nearly identical model neurons that do not employ response normalization. Their response drives are equal to **f**^*T*^**c** instead of **f**^*T*^**c**/*N*. The neurons prefer the same feature and differ only because one has a mismatched and the other has a matched weight matrix. The larger mismatched matrix is padded with zero-valued coefficients. Multiplying inputs with zero-valued coefficients does not change the linear response. Both neurons will thus produce identical linear responses regardless of whether the matrix is matched. The differential effects must therefore be due to the size of the image region that determines the normalization factor, relative to the size of the preferred feature of the receptive field.

### Downsampling

The model neuron receptive fields considered thus far have had identical sampling resolution, so the number of pixels representing a preferred feature scales with the visual angle spanned by the receptive field (Fig. 9A,B). In the primate visual system, at a given eccentricity, simple cells with larger receptive fields pool over more relay cell inputs from the lateral geniculate nucleus (LGN) than those with smaller receptive fields (Taylor, Sedigh-Sarvestani, Vigeland, Palmer, & Contreras, 2018). Similarly, parasol ganglion cells in the retina pool over more cone receptors than midget ganglion cells (G. D. Field et al., 2010). Sometimes, however, large receptive fields pool inputs that have lower sampling resolution than their smaller counterparts. For example, large retinal ganglion cells (RGCs) in the retinal periphery pool inputs from large low-resolution cone photoreceptors, whereas small foveal RGCs of the same type pool over small high-resolution photoreceptors (Croner & Kaplan, 1995; Rossi & Roorda, 2010). The processing motif employed by the peripheral retina is roughly equivalent to downsampling, a common pre-processing method in the computer vision, image processing, and deep learning communities (Burt & Adelson, 1983). In general, downsampling reduces the number of pixels (i.e. sampling resolution) representing a particular image patch, and hence the computational requirements for processing that patch.

**Figure 9.**
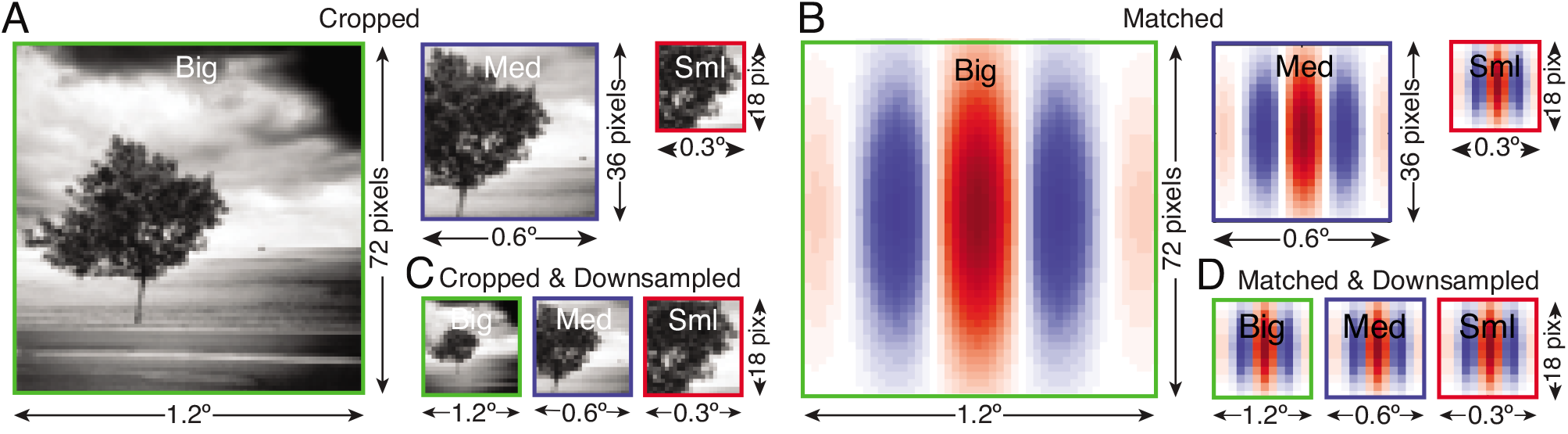
Downsampling cropped images for matched receptive field weight matrices. **A** Cropped images: big, medium, and small. Boxes indicate three different scales at which image data is processed. The cropped images span different visual angles but have a fixed sampling rate. Each cropped image therefore has a different number of pixels. **B** Matched receptive field weight matrices. The visual angle spanned by the weight matrix matches the visual angle spanned by the preferred feature. Each matrix also has a different number of pixels. **C** Cropped and downsampled images. Cropped and downsampled images span different visual angles, but have the same number of pixels. **D** Matched and downsampled receptive fields. All weight matrices have the same number of pixels, regardless of the spanned visual angle.

**Figure 10.**
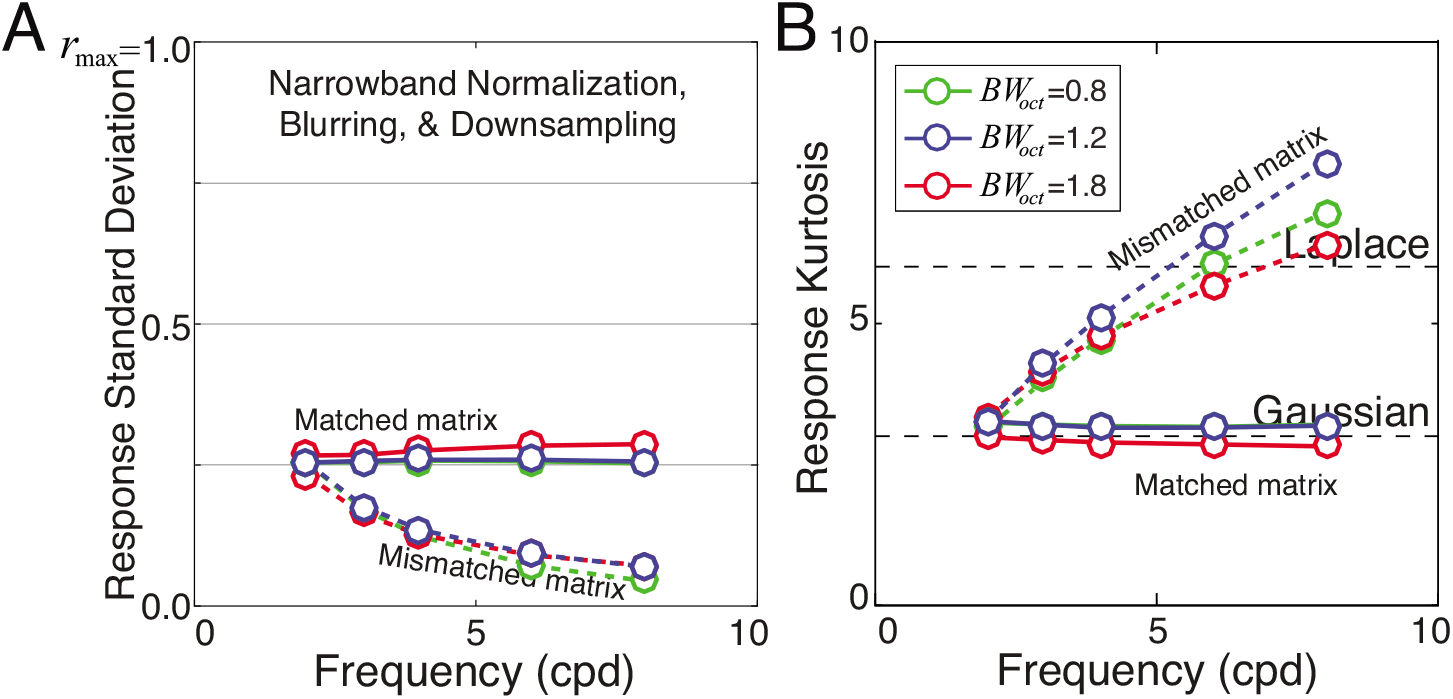
The impact of downsampling on response statistics. **A** Stimulus-driven response standard deviation with narrowband normalization, blurring, and downsampling. **B** Stimulus-driven response kurtosis with blurring and downsampling. The response statistics are essentially identical with and without downsampling.

We asked how downsampling the input stimuli impacts model neuron response statistics. First, we generated a new set of receptive field weight matrices where both the spanned visual angle and the sampling resolution were yoked to the preferred feature. The result was a set of receptive fields defined by weight matrices that all had an identical number of pixels. Specifically, all weight matrices had 18×18 pixels, the same number as the original matrix corresponding for the smallest preferred feature (0.3°, 8 cpd). Then, we downsampled the image patches (after appropriate blurring to prevent aliasing) to match the sampling resolution of the receptive fields (see Methods). Downsampled images and receptive fields are shown in Figure 9C,D.

With downsampling, the narrowband-normalized responses with matched matrices are constant variance Gaussian (Fig. 10A,B). The response statistics (i.e. standard deviation and kurtosis) are within one percent of the response statistics without downsampling (see Fig. 7DE). There is no disadvantage (or advantage) to downsampling in terms of the signal-to-noise for signal discrimination. Thus, at least in terms of signal-to-noise for stimulus discrimination, there is no pressure on the visual system to avoid downsampling. This result is useful for approaches that seek to learn receptive fields via non-parametric methods, where every additional pixel in a receptive field weight matrix incurs considerable computational cost (see Discussion).

## Discussion

Model neurons employing narrowband response normalization yield scale-invariant Gaussian distributed responses to natural stimuli. The scale-invariant response statistics come close to maximizing the signal-to-noise ratio for stimulus discrimination with natural stimuli, but the scale-invariance depends on the normalization factor being determined from image region that matches the size of the receptive field’s preferred feature. In the discussion section, we examine how these results are affected by receptive fields that are not oriented Gabors, and stimulus types that are not natural (i.e. noise stimuli). We discuss how the results reported here can explain why models fitted to neurons in cortex tend to poorly predict responses to natural stimuli, even when they beautifully predict responses to noise stimuli.

### Generality of conclusions

Different subfields in vision and computational neuroscience have different methodological conventions for modeling neurons. Under many simplified circumstances, the different conventions have little or no practical impact. However, when the model neurons include the dominant features of real neurons in cortex—receptive field, response normalization, and encoding noise—the subtle differences in the modeling conventions can have a dramatic impact on their response statistics to natural images. How, then, do modeling choices other than those considered in the body of the paper impact the response statistics?

First, we asked whether the response statistics generalize to other Gabor receptive fields. In the results section, we analyzed only the response statistics of vertically oriented even-symmetric (cosine phase) Gabor receptive fields. Do odd-symmetric (sine phase) receptive fields produce similar results? Yes. Gabor receptive fields with all phases and orientations produce equivalent results. Thus, model neurons with biologically plausible receptive fields and appropriate narrowband response normalization produce response statistics that are invariant to the preferred feature.

Next, we asked whether model neurons with other receptive field shapes produce similar results. There is a long history of using Gabor functions to describe the receptive fields of simple cells in early visual cortex (Jones & Palmer, 1987a; 1987b). However, empirical data suggests that log-Gabors may provide a better characterization of simple cell receptive fields in early visual cortex (De Valois, Albrecht, & Thorell, 1982a; Geisler & Albrecht, 1997; Hawken & Parker, 1987). It has also been argued on theoretical grounds that log-Gabor receptive fields may be better than Gabor receptive fields for encoding natural images (D. J. Field, 1987). We re-ran our analyses with log-Gabor shaped receptive fields. All results hold with log-Gabor receptive fields (Fig. S7).

Then, we asked whether the main conclusions hold for the receptive fields like those of retinal ganglion cells or relay cells in the lateral geniculate nucleus, which are radially symmetric and do not select for orientation. We repeated our analyses with center-surround Gabors and difference-of-Gaussian (DoG) preferred features embedded in matched weight matrices. Results are quite similar for oriented and un-oriented receptive fields with narrowband normalization. However, model neurons with un-oriented receptive fields also yield Gaussian response distributions with broadband normalization (Fig. S8). (Recall that oriented receptive fields with broadband normalization yield Laplace-distributed responses; Fig. 4C,D, Fig. S2) This result implies that the differences in the shape of the response distributions that distinguish broadband and narrowband normalization with oriented Gabors (c.f. Fig. 4) are due primarily to the orientation selectivity of the receptive field.

### Natural vs. noise stimuli

Model neurons with narrowband response normalization and matched weight matrices yield Gaussian-distributed responses to natural stimuli. But noise stimuli are more often used in psychophysical and neurophysiological experiments because they have useful properties for methods designed to recover the stimulus features (i.e. receptive fields) that drive behavioral and neural response (Ahumada & Lovell, 1971; Schwartz, Pillow, Rust, & Simoncelli, 2006). 1/f noise has the amplitude spectrum but not the phase structure of natural images. White noise has neither the amplitude spectrum nor the phase structure of natural stimuli. It is therefore important to ask how our model neurons respond to noise stimuli.

We compared linear and normalized model neuron responses to natural, 1/f noise, and white noise stimuli. As shown above, linear responses to natural stimuli are distributed with heavier tails than Laplace distributions. Broadband-normalized responses to natural stimuli are Laplace-distributed. And narrowband-normalized responses to natural stimuli are approximately Gaussian (Figs. S1-3). In contrast, linear and broadband responses to Gaussian 1/f and white noise stimuli are Gaussian. Narrowband-normalized responses to 1/f and white noise stimuli are also approximately Gaussian (Figs. S9,S10). Thus, the differences between linear, broadband-, and narrowband-normalized responses are substantial for natural stimuli. The negligible for noise stimuli (Fig. 11).

**Figure 11.**
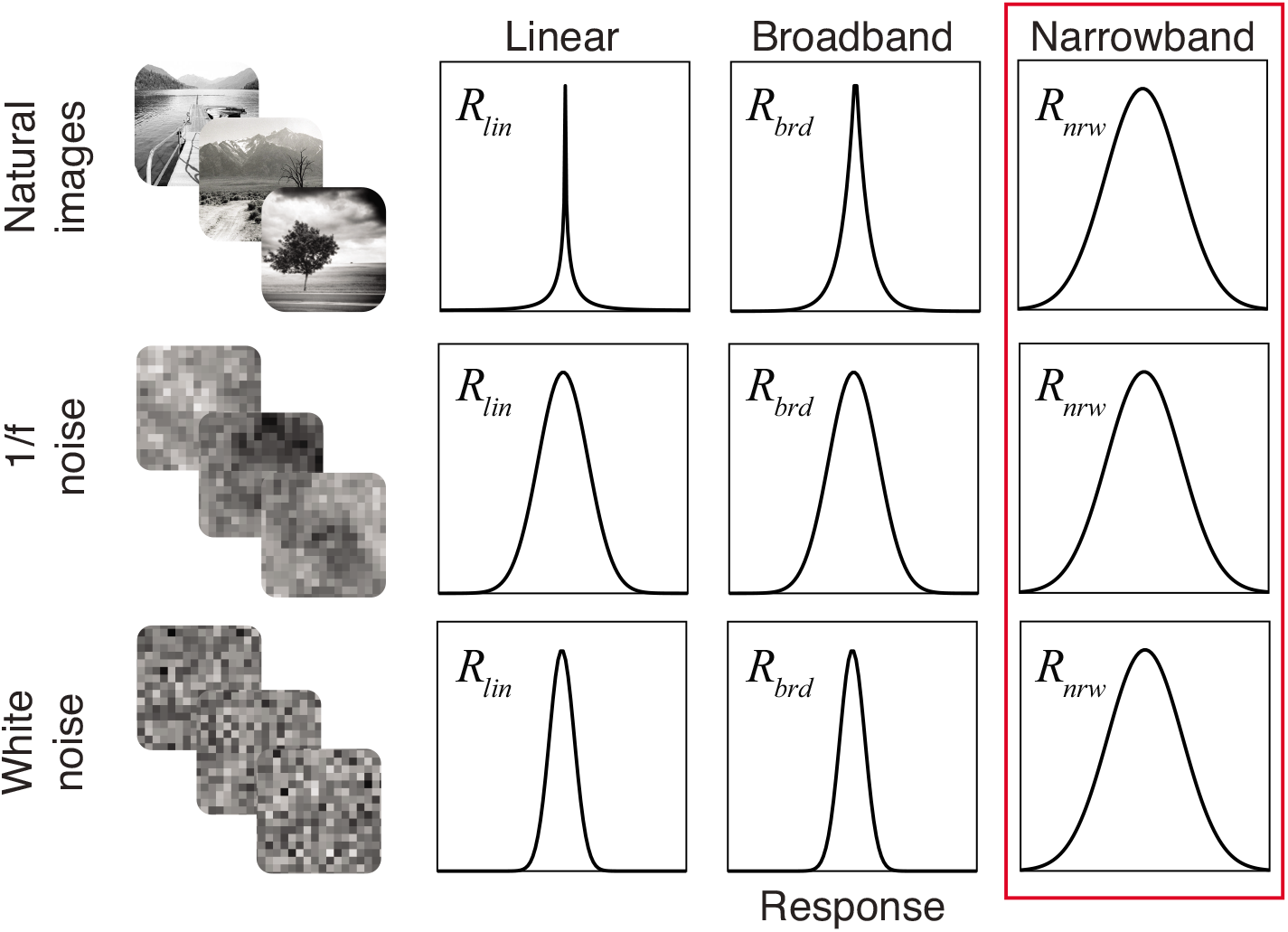
Distributions of linear, broadband-normalized, and narrowband-normalized responses to natural, 1/f noise, and white noise stimuli. Linear responses to natural stimuli are very heavy tailed (i.e. heavier than a Laplace distribution; see Fig. S1). Broadband responses to natural stimuli are Laplace-distributed (see Fig. S2). Narrowband responses to natural stimuli are Gaussian (see Figs. 4, 7, S3). Linear and broadband responses to 1/f and white noise are guaranteed to be Gaussian, assuming that the number of pixels defining each stimulus (i.e. dimensionality) is high enough. Narrowband responses to noise stimuli are all approximately Gaussian. Narrowband normalization produces responses (i.e. response drives) that are very nearly Gaussian with all three stimulus types. Narrowband normalization helps standardize the distributional form of the response statistics across stimulus types (box). Narrowband normalization should therefore improve the ability of computational models of visual information processing to generalize across stimulus types.

These findings provide a clue about why subunit models of neural response tend to generalize poorly when tested with natural stimuli. Subunit models are a popular method for performing neural systems identification. Their aim is provide a concise computational level description of the input-output relationship between stimulus and response. Subunit models are typically fit and tested using noise stimuli. But even when fitted subunit models nicely predict performance with noise stimuli, they tend to perform poorly when tested with natural stimuli (Heitman et al., 2016; Smyth, Willmore, Baker, Thompson, & Tolhurst, 2003; Talebi & Baker, 2012).

Subunit models of neural response tacitly assume Gaussian-distributed responses (McFarland, Cui, & Butts, 2013; I. M. Park, Archer, Priebe, & Pillow, 2013; Rust, Schwartz, Movshon, & Simoncelli, 2005; Schwartz et al., 2006), even though the models do not typically include response normalization (but see (McFarland et al., 2013)). However, with Gaussian noise stimuli linear (un-normalized) response distributions are guaranteed to be Gaussian. (Broadband-normalized response distributions to noise stimuli are also guaranteed to be Gaussian, assuming that the number of pixels defining each stimulus—i.e. the dimensionality of each stimulus—is sufficiently high (Poincaré, 1912).) Thus, the absence of narrowband response normalization has little practical effect with the stimuli that subunit models are most often fit and tested. In contrast, with natural stimuli, the absence of narrowband response normalization results in response distributions that are highly non-Gaussian. The non-Gaussian response distributions violate the tacit distributional assumptions of the models. This violation contributes to the poor generalization of subunit models to natural stimuli. Incorporating narrowband normalization into these methods for neural systems identification will yield Gaussian response distributions for both natural and noise stimuli, and should therefore improve the ability of these models to generalize across these stimulus types.

### Non-parametric receptive field learning: Model neuron modeling conventions

Some areas of computational neuroscience aim to learn populations of receptive fields (i.e. preferred features) that optimize a particular goal. The preferred features of these receptive fields are typically learned by iteratively updating coefficients (i.e. pixel values) in pursuit of the goal. The receptive field weight matrices typically have a fixed number of pixels and span a fixed visual angle. This convention is convenient for matrix-based programming languages (e.g. Matlab), but it often results in weight matrices that are mismatched to the preferred feature. For example, mismatched matrices are commonly reported by papers that fit subunit models to describe the stimulus-response properties of neurons early in the visual processing stream (c.f. Fig. 2a in Rust et al, 2005; (McFarland et al., 2013; I. M. Park et al., 2013; Samengo & Gollisch, 2013; Schwartz et al., 2006; Vintch, Movshon, & Simoncelli, 2015)). Analyses inspired by the efficient coding hypothesis seek receptive field populations that efficiently encode natural stimuli also commonly report mismatched weight matrices (c.f. Fig. 4a in Olshausen & Field, 1996; (Bell & Sejnowski, 1997; Lewicki, 2002; Olshausen & Field, 1997; Rehn & Sommer, 2007)). Why are mismatched weight matrices commonly reported if they carry the disadvantages detailed in the results section (c.f. Fig. 8)? The literatures mentioned above typically assume that receptive field (or subunit) responses are driven purely by linear operations; that is, they do not incorporate response normalization. (They also do not typically model encoding noise.) In the absence of normalization, matched and mismatched matrices yield identical responses (see Results).

To increase biological realism and to facilitate generalization to natural stimuli, normalization should be included in the response models for these feature-learning methods and others. When normalization is included, the normalization factor must be computed from the same stimulus region as the preferred feature. Otherwise, receptive fields with preferred features smaller than the visual angle spanned by the weight matrix will have poor signal-to-noise (Fig. 7,8). This will make it difficult to interpret the results of feature learning studies. If high frequency features are not learned, it will be unclear whether it is because of the poor signal-to-noise or because high frequency filters are fundamentally not useful. If receptive fields within a parametric family are being learned (e.g. Gabor) the parameters can be used to determine the area spanned by the feature, but this is not possible with non-parametric approaches. One way to address this issue would be learn receptive fields with localized priors, but this technique poses significant technical challenges in many contexts (M. Park & Pillow, 2011). A simpler approach would be to learn receptive fields with response normalization at multiple scales simultaneously. At each scale, receptive fields with small mismatched preferred features will yield with poor signal-to-noise, biasing the learning procedures against selecting those features relative to the scale. But across scales, large and small features will both be given a fair chance; large features would be matched to large matrices and small features would be matched to small matrices.

### Neural response in early visual cortex

Across many natural images, model neuron response drive is mean zero and Gaussian distributed. The response drive is equivalent to the neural response under the assumptions that i) no response rectification occurs and ii) that the power of the static output non-linearity is 1.0. In real neurons these assumptions do not typically hold. As a consequence, the responses of real neurons in early visual to natural stimuli are not Gaussian (Felsen et al., 2005; Weliky et al., 2003). Rather, the response drive is typically rectified and the power of static output nonlinearity is close to 2.0 (Priebe & Ferster, 2008). We examined how rectification and a squaring output non-linearity changes the model neuron response statistics. Rectifying and squaring the response drive converts the Gaussian response distributions into chi-squared response distributions with one degree of freedom 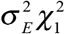 scaled by the stimulus-driven response variance (Fig. 12). The scaled chi-squared is more similar to the response distributions observed from spiking neurons in cortex. The scaled chi-squared distribution is equivalently described by a gamma distribution with shape and scale parameters of 1/2 and 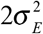, respectively.

These considerations make several predictions. First, presenting natural stimuli to simple cells in early visual cortex should yield response rate distributions that are nicely modeled by scaled chi-squared distributions with one degree of freedom. Second, because complex cells are typically modeled as resulting from quadratic pooling of two (or more) subunit receptive field responses, complex cells should yield response rate distributions that are well modeled by scaled chi-squared distributions with two (or more) degrees of freedom. Third, the distribution of response across stimuli for different neurons should be approximately invariant to the preferred feature of each neuron. We advocate examining the distribution of response across hundreds (or thousands) of natural images and comparing their distribution to a chi-squared distribution with variable degrees of freedom.

**Figure 12.**
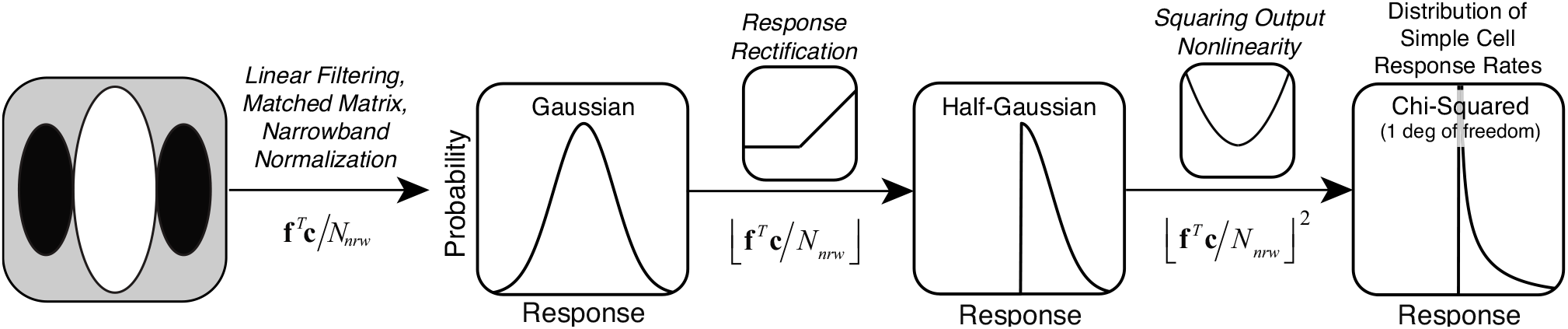
Relating Gaussian response drive statistics to response statistics in cortex. Gaussian responses are first rectified to create a half-Gaussian distribution. Rectified responses are then squared to create a scaled chi-squared distribution (see text). The scaled chi-squared distribution represents the predicted distribution of response rates to natural images by neurons (i.e. simple cells) in early visual cortex.

It is also important to ask how the signal-to-noise ratio for stimulus discrimination would be impacted by rectifying and squaring the response drive (i.e. the normalized linear response). If the controlling source of encoding noise occurs after rectification and squaring, signal-to-noise would be altered (Eq. 7A). On the other hand, if the controlling source of encoding noise is added at the level of the response drive, before the rectification and squaring (Eq. 7B), the signal-to-noise for stimulus discrimination will be unaffected because a monotonic transform of a noisy signal together will not alter the signal-to-noise ratio (Pelli, 1985; Rieke & Rudd, 2009).

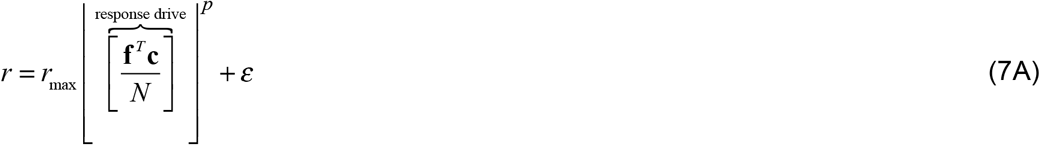

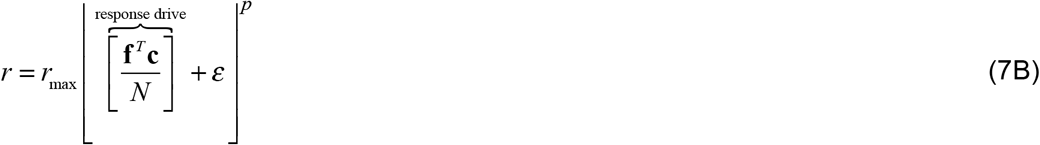

A number of influential models in the neurophysiological literature propose that the controlling source of encoding noise is at the level of the membrane voltage, and should be modeled as constant, additive, and zero mean (Carandini, 2004; Mohanty, Scholl, & Priebe, 2012; Priebe & Ferster, 2008). After rectification and squaring, the noisy voltage signal predicts Poisson-like encoding noise (i.e. response variance that scales with the mean response to each stimulus) like that typically observed in cortex (Tolhurst, Movshon, & Dean, 1983b). If these models are correct, the controlling encoding noise should modeled as occurring at the level of the response drive (Eq. 7B). Lastly, these considerations make one an additional prediction. If the intracellular voltage reflects the response drive as it has been discussed in this manuscript, the distribution of intracellular voltages across the natural stimulus ensemble should be Gaussian distributed.

### Task specific analysis of natural images and scenes

Efficient coding is an influential theoretical framework for thinking about how neural systems encode natural stimuli (Attneave, 1954; Barlow, 2001). Many papers have focused on learning receptive fields that efficiently reconstruct proximal stimuli (Bell & Sejnowski, 1997; Lewicki, 2002; Olshausen & Field, 1996; 1997). However, the sensory-perceptual systems of humans and other animals must do more than efficiently encode and reconstruct sensory inputs. Sensory-perceptual systems must extract information from input stimuli that is useful for the behavioral tasks that organisms must perform to survive and reproduce. Efficient coding does not directly address this problem.

In recent years, statistical techniques have been developed that learn small populations of receptive fields that encode the stimulus features most useful for specific tasks (Burge & Jaini, 2017; Geisler, Najemnik, & Ing, 2009; Jaini & Burge, 2017). These techniques have helped to find the optimal solutions in tasks useful for estimating the three-dimensional structure of the environment—focus error estimation (Burge & Geisler, 2011; 2012), disparity estimation (Burge & Geisler, 2014), and retinal motion estimation (Burge & Geisler, 2015)—and that have been the focus of intense study for decades by the vision and neuroscience communities (Banks, Gepshtein, & Landy, 2004; Burge, Fowlkes, & Banks, 2010; Burge, Peterson, & Palmer, 2005; Cormack, Stevenson, & Schor, 1991; Flitcroft, 1990; Gekas, Meso, Masson, & Mamassian, 2017; Held, Cooper, & Banks, 2012; Iyer & Burge, 2018; Jogan & Stocker, 2015; Kotulak & Schor, 1986; Kruger, Aggarwala, Bean, & Mathews, 1997; Priebe & Lisberger, 2004; Priebe, Cassanello, & Lisberger, 2003; Rust, Mante, Simoncelli, & Movshon, 2006; Sebastian, Burge, & Geisler, 2015; Simoncelli & Heeger, 1998; Tyler & Julesz, 1978; Weiss, Simoncelli, & Adelson, 2002). In this manuscript, we examined only the response statistics of individual model neurons and considered only performance in a very simple task: discriminating one stimulus from another. The optimal solutions to these more sophisticated tasks require combining the responses from multiple receptive fields. With appropriate normalization, responses to natural stimuli from multiple receptive fields are jointly Gaussian, a fact that should simplify computations for optimally combining those receptive field responses. Thus, the results reported here should be thought of as the beginning of a more complete investigation of how visual systems process natural stimuli. Having an accurate picture of the response statistics of model neurons, and understanding how small differences in modeling conventions affect those response statistics, lay an important foundation for building principled models in the future.

## Conclusion

The work presented in this manuscript suggests that computational models of neural processing should incorporate narrowband (i.e. stimulus- and feature-specific) response normalization. A simple expression for computing the narrowband normalization factor is provided, that should facilitate the inclusion of narrowband normalization in non-parametric methods for learning receptive fields of model-neuron-like units. Narrowband normalization should also improve the ability of such models to generalize from noise stimuli to natural stimuli, because narrowband normalization yields scale invariant Gaussian responses for both natural and noise stimuli.

## Acknowledgments

This work was supported by startup funds to JB from the University of Pennsylvania, and by NIH grant R01-EY028571 to JB from the National Eye Institute and the Office of Behavioral and Social Sciences Research. We thank Benjamin Chin for helpful comments on a draft version of the manuscript and Takahiro Doi for helpful discussions.

## Methods

### Natural stimuli

Natural image patches were sampled from two recently published photographic databases of natural scenes (Burge & Geisler, 2011; Burge, McCann, & Geisler, 2016). Scenes were photographed on and around the University of Texas at Austin campus and contained grass, shrubs, and trees, and streets, cars, and buildings. The images were calibrated such that the intensity values were linear with luminance. The data represents 30,888 unique 1.2° image patches (72×72 pixels); 312 non-overlapping patches were randomly selected from each of 99 calibrated natural images.

### Local contrast

The intensity patches are converted to Weber contrast images by luminance normalization. The contrast image is obtained by subtracting off and dividing by the local mean intensity

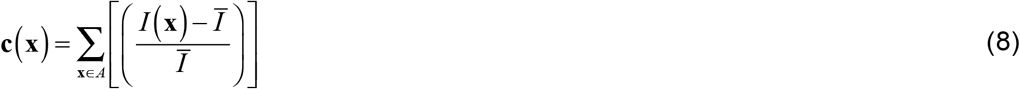

where **c**(**x**) is the local contrast image patch, *I*(**x**) is the local intensity image patch, ***I̅*** is the local mean intensity, and **x** = {x,y} indexes spatial position in the area *A* spanned receptive field weight matrix. The local mean intensity is given by 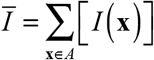.

### Receptive fields

The receptive field of each model neuron is modeled with a weight matrix. The receptive field weight matrix is determined by the preferred feature, the visual angle spanned by the weight matrix, and by the spatial sampling rate. The preferred feature of each model neuron is modeled as a Gabor. A Gabor is a cosine wave multiplied by a Gaussian envelope

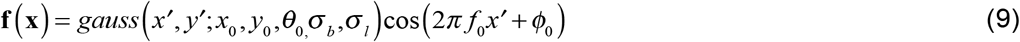

where *x*_0_ and *y*_0_ specify the position of the Gaussian envelope, *θ*_0_ is the preferred orientation, *σ_b_* is the standard deviation of the envelope in the band pass direction (orthogonal to the grating orientation), *σ_l_* is the standard deviation of the envelope in the low pass direction (parallel to the grating orientation), *f*_0_ is the preferred spatial frequency, *φ*_0_ is the preferred phase, and {*x*′,*y*′} are transformed coordinates due to the preferred orientation

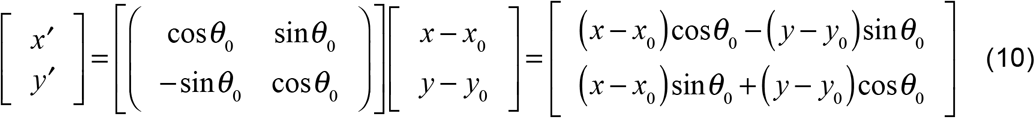

The coefficients of the receptive field weight matrix are normalized such that the L2 norm of the receptive field weight coefficients 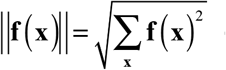 equals 1.0. The octave bandwidth of the preferred feature is given by the log-base-two ratio of the high and low frequencies at half-height

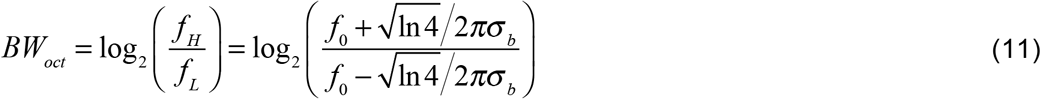

The orientation bandwidth specifies the polar angle spanned by the Gaussian envelope at halfheight and is given by

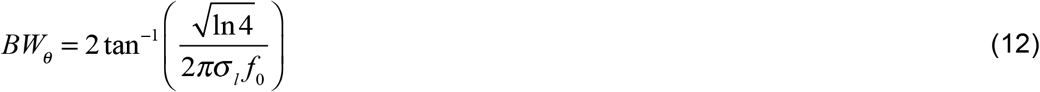

When the visual angles spanned by the *x*′ and *y*′ values are 5x the envelope standard deviations in the band pass and low pass directions, respectively, the weight matrix is matched to the preferred feature. When the spanned visual angles are greater than 5x the standard deviation in either the band pass or low pass direction, the weight matrix is mismatched to the receptive field.

We analyze the response statistics of model neurons with vertically-oriented Gabor receptive fields having 42° orientation bandwidths and 0.8, 1.2, and 1.8 octave bandwidths. Simple cells in early visual cortex have a median orientation bandwidth of 42°, a median octave bandwidth of 1.2 octaves. The distribution of cortical octave bandwidths spans approximately 0.8 to 1.8 octaves at half-height (De Valois, Albrecht, & Thorell, 1982a; De Valois, Yund, & Hepler, 1982b). We computed response statistics for mismatched receptive field weight matrices spanning 5x (e.g. 2 cpd, 72 pixels, 1.2°) to 20x (e.g. 8 cpd, 72 pixels, 1.2°) the envelope standard deviations.

The aspect ratio of the Gaussian envelope in terms of the octave and orientation bandwidths is obtained by solving Eqs. 11 and 12 for *σ_b_* and *σ*_l_, respectively, and then taking the ratio

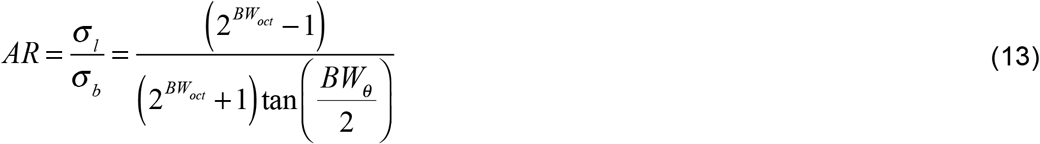

The log-base-two aspect ratios log_2_(*AR*) of these receptive fields are −0.5, 0.0, and 0.5, respectively, which correspond to envelopes that are wider than high by a factor of 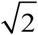, circular, and higher than wide by a factor of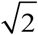, respectively.

All data in the main text is presented for rectangular image patches and receptive field weight matrices. However, there are practical disadvantages to working with matrices that are rectangular. It is more convenient to work with square patches and weight matrices. We examined how the response statistics differ between rectangular and square weight matrices. Note that nominally matched square matrices are actually slightly mismatched for octave bandwidths other than 1.2. We computed the response statistics with square image patches and weight matrices for all octave bandwidths. The differences were minor (Fig. S11).

### Normalization

To obtain the normalization factor for each stimulus, we converted each contrast image into its frequency domain representation by performing a fast-fourier transform (FFT). Next, we normalized the transform such that its total power equaled the total energy of the contrast image, in accordance with Parseval’s theorem. To prevent high-frequency artifacts that may be caused by the edge of the image patch, it is common to apply a cosine window before performing the FFT. However, because of numerical issues, it is impossible to avoid occasionally exceeding the maximum response *r*_max_ when a window is applied. Stimulus energy near the edge of the image patch that increases the linear response, may be windowed out of the normalization factor. In this case, the normalization factor will be smaller than it should be. In some cases, it will cause the normalized response to exceed the maximum. Results were similar with and without windowing, but they were better behaved without windowing.

### Encoding noise

The responses of neurons are noisy. If the same exact stimulus is presented multiple times, the neuron is likely to give a slightly different response to each presentation. We considered two types of encoding noise: constant additive noise and scaled additive noise. Both types were modeled as mean zero Gaussian noise 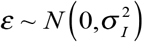. With constant additive noise, the encoding noise variance 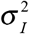 is constant regardless of the mean response. With scaled additive noise, the encoding noise variance 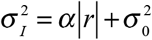 scales in rough proportion to the mean, where *α* is the fano factor. All of the qualitative results are essentially invariant to whether constant or scaled additive noise is used. The manuscript presents results for constant additive noise.

### Downsampling

Image patches were downsampled using Matlab’s imresize.m function with linear interpolation. Similar results are obtained using Matlab’s impyramid.m. However, with impyramid.m the downsampling factors are restricted to powers of two; we favor imresize.m because of its increased flexibility. Other downsampling methods are likely to produce very similar results.

## Supplement

**Figure S1.**
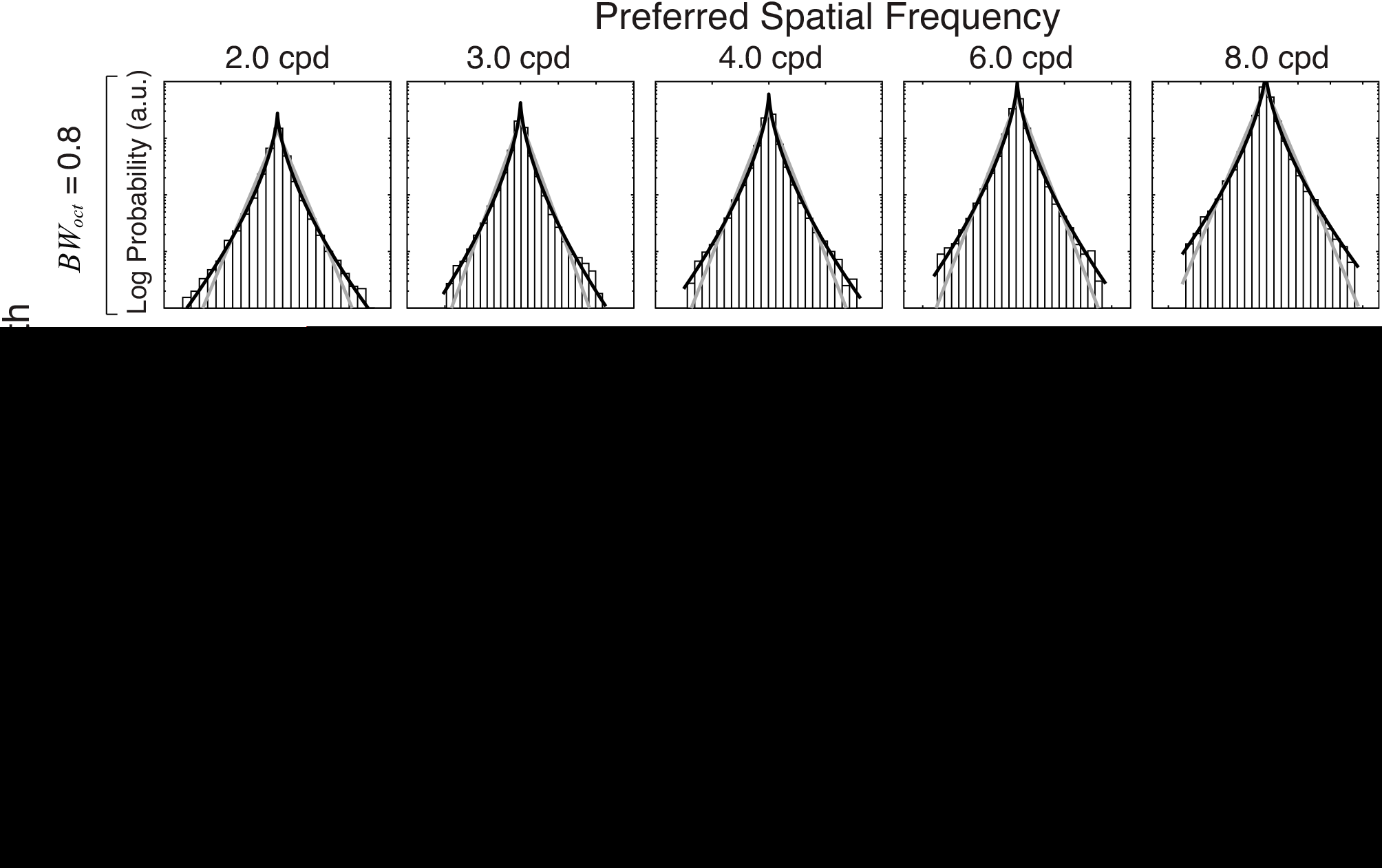
Linear receptive field responses to natural images for all preferred spatial frequencies (columns) and octave bandwidths (rows). Note the dramatic difference in response magnitude (x-axis) as a function of preferred frequency. The responses are fit with both a Laplace and a generalized Gaussian via maximum likelihood (gray & black curves, respectively). The best-fit generalized Gaussian fit has significantly heavier tails than the best fit Laplace in all cases. The generalized Gaussian is given by 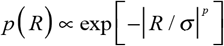 where the Laplace distribution (*p* =1.0) and the Gaussian (*p* =2.0) are special cases. The linear responses are best fit with powers *p* of between 0.62 and 0.70, with a mean power of 0.65. The y-axis indicates log-probability over a four order-of-magnitude range. The Laplace fit appears as straight lines on a log-probability plot.

**Figure S2.**
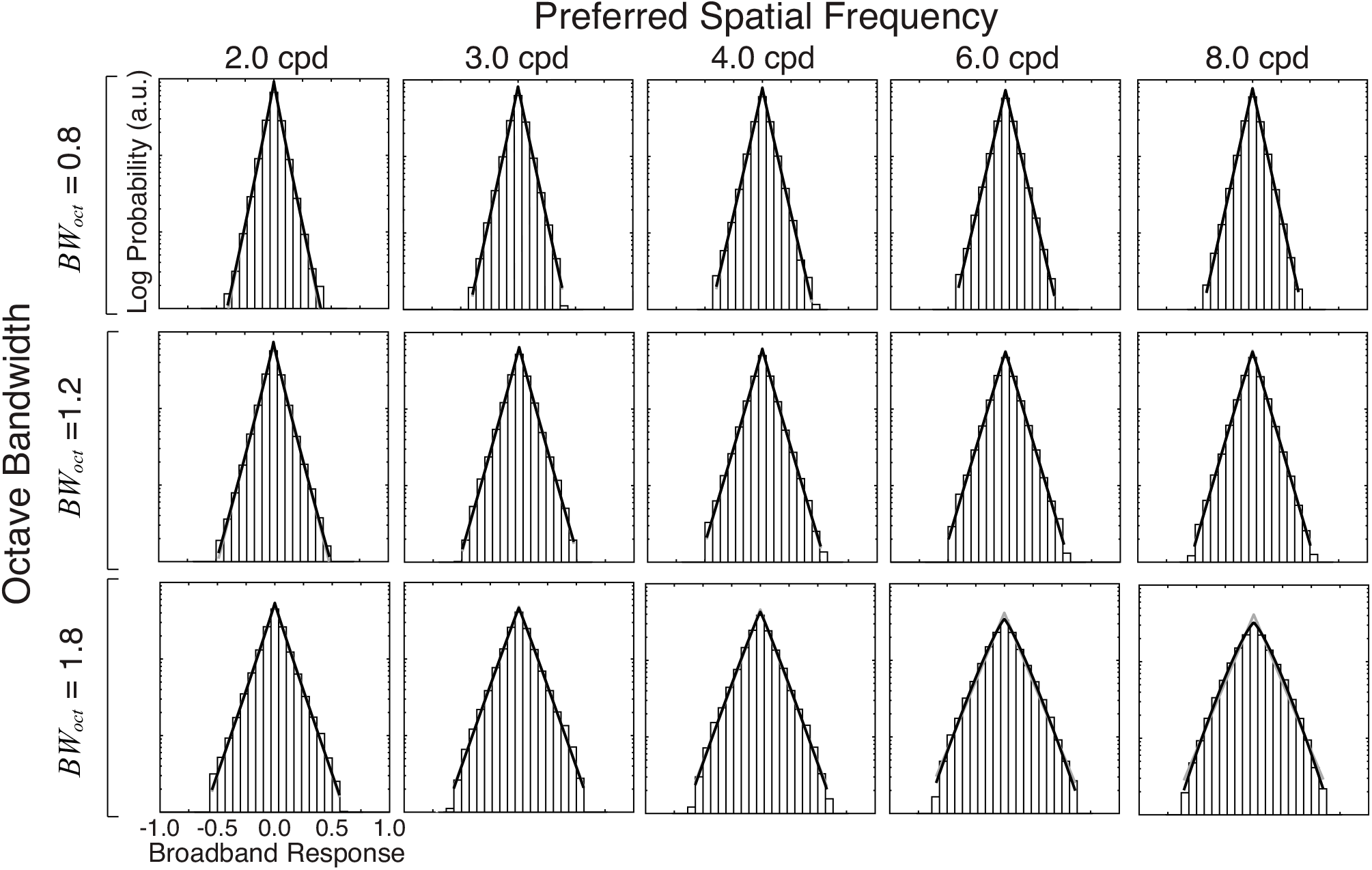
Broadband-normalized responses to natural images are Laplace distributed. Broadband responses with matched receptive field weight matrices for all preferred spatial frequencies (columns) and octave bandwidths (rows). The responses are fit with both a Laplace and a generalized Gaussian via maximum likelihood (gray & black curves, respectively). The generalized Gaussian is given by 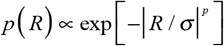 where the Laplace distribution (*p* =1.0) and the Gaussian (*p* =2.0) are special cases. The generalized Gaussian fit is indistinguishable from the Laplace fit in almost all cases; when the gray lines are not visible, it is because they are lying behind the black curve. The y-axis indicates log-probability over a three order-of-magnitude range. The Laplace fit appears as straight lines on a log-probability plot.

**Figure S3.**
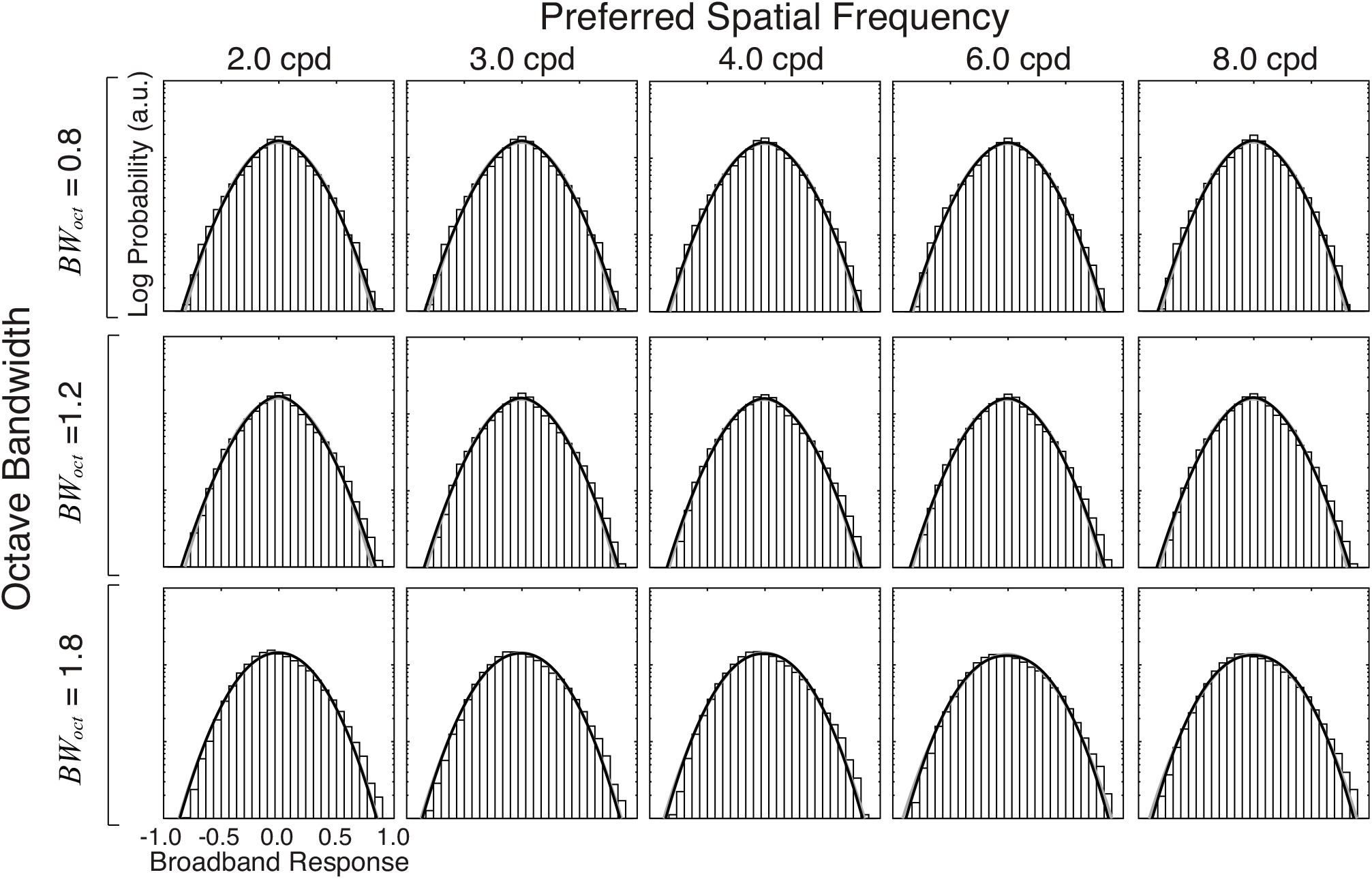
Narrowband-normalized responses to natural images are Gaussian-distributed. Narrowband responses with matched receptive field weight matrices for all preferred spatial frequencies (columns) and octave bandwidths (rows). The responses are fit with both a Gaussian and a generalized Gaussian via maximum likelihood (gray & black curves, respectively). The generalized Gaussian is given by 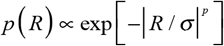 where the Laplace distribution (*p* =1.0) and the Gaussian (*p* =2.0) are special cases. The generalized Gaussian fit is indistinguishable from the Gaussian fit in almost all cases; when the gray lines are not visible, it is because they are lying behind the black curve. The y-axis indicates log-probability over a three order-of-magnitude range. The Gaussian distribution appears as an upside-down parabola on a log-probability plot.

**Figure S4.**
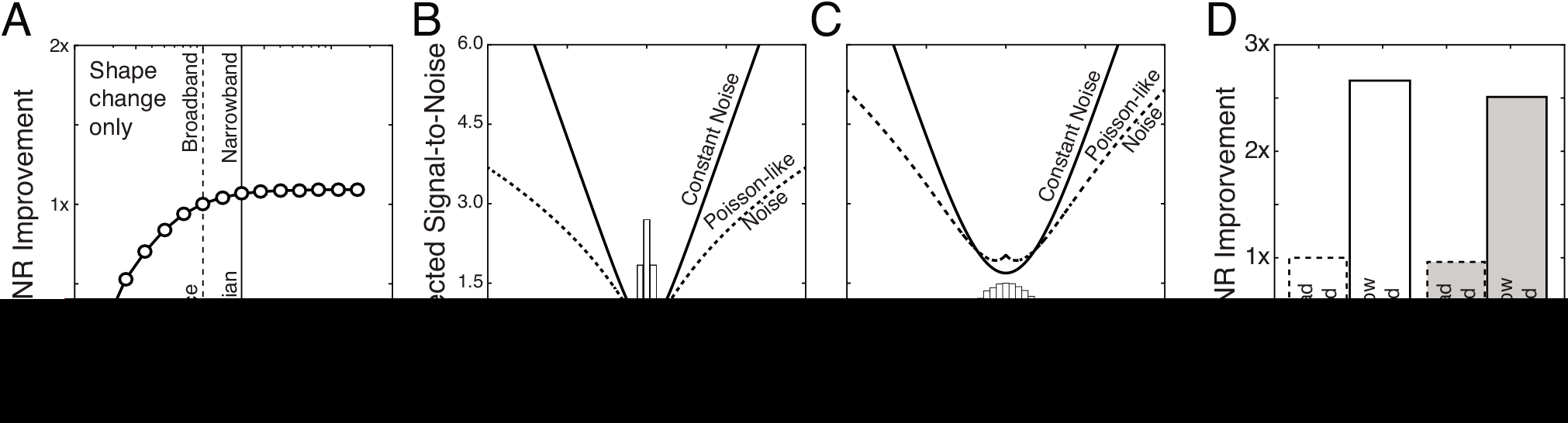
The impact of the shapes of the response distributions and the type of encoding noise on SNR. **A** The impact of the shape of the response distribution on signal-to-noise ratio for stimulus discriminability, assuming constant encoding noise. We computed expected SNR for a set of simulated generalized Gaussian response distributions having identical means and variances but different powers. The generalized Gaussian is given by 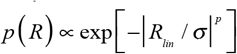. The Laplace distribution (*p* =1.0; kurtosis = 6.0) and the Gaussian (*p* =2.0; kurtosis = 3.0) are special cases. Powers larger than 2.0 approach the uniform distribution. **B** Expected SNR for broadband-normalized responses as a function of the response for constant (solid curve) and Poisson-like (dashed curve) encoding noise. The histogram shows the distribution of stimulus-driven broadband response. The constant Gaussian encoding noise was set such that the expected SNR from the broadband responses across the stimulus ensemble was equal to 1.0 (i.e. *σ_I_* ≅ *σ_E_*/3). The Poisson-like is encoding noise is given by 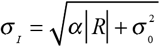 where *α* is the fano factor and 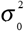 are the fano factor and baseline variance, respectively. The values of the fano factor and the baseline variance were taken from the neurophysiological literature. Finally, the mean Poisson-like encoding noise variance across the stimulus ensemble was matched to the constant encoding noise variance. SNR for stimulus discrimination is approximately 4% lower with Poisson-like encoding noise. **C** Expected SNR for narrowband-normalized responses as a function of the response. The mean encoding noise variance was matched to the mean encoding noise for the broadband responses. SNR for stimulus discrimination is approximately 8% lower with Poisson-like encoding noise. **D** Summary of expected SNR improvement for broadband and narrowband normalization, with constant and Poisson-like encoding noise.

**Figure S5.**
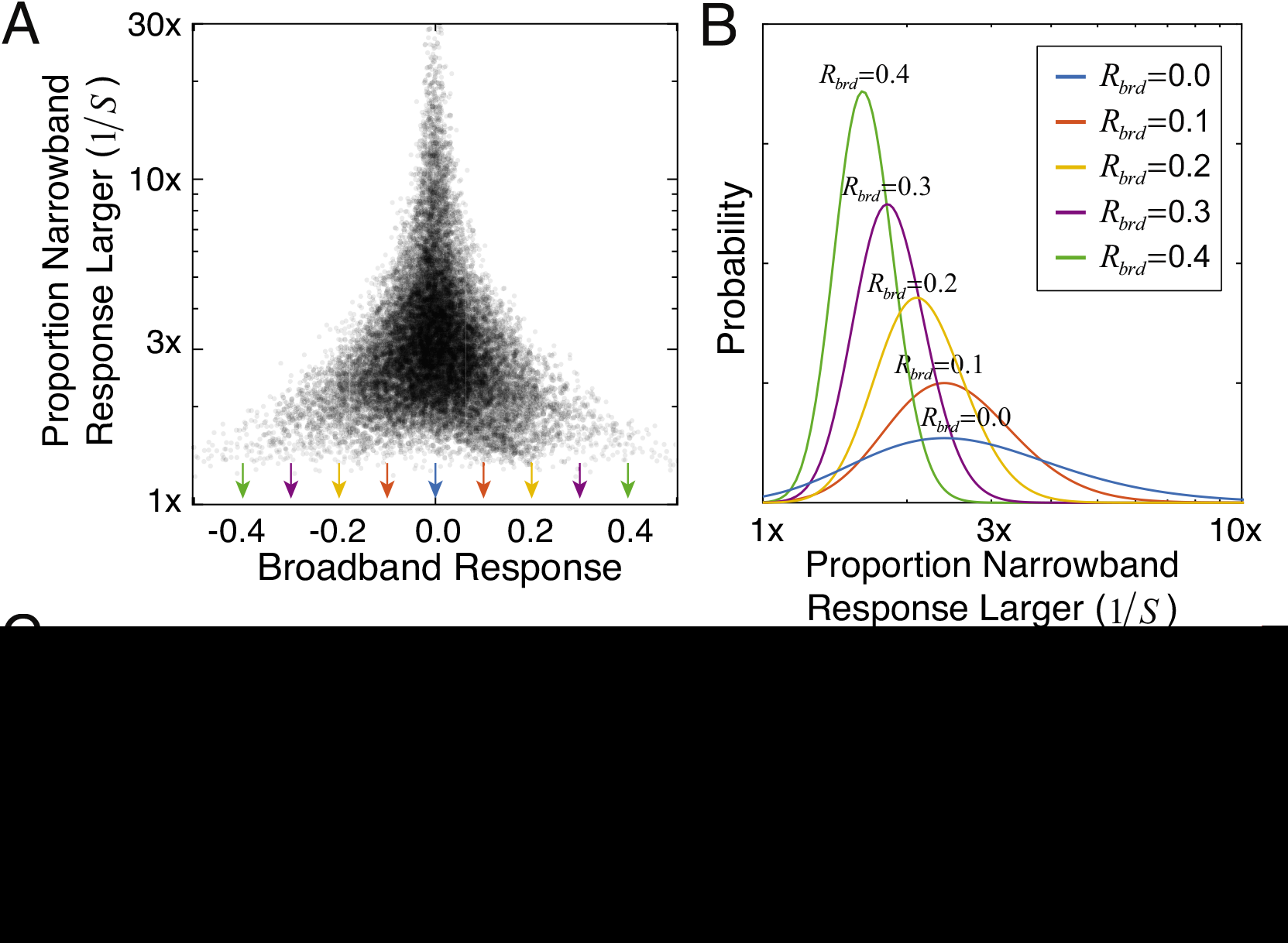
The impact of narrowband normalization. **A** Proportion that the narrowband response is larger than the broadband response *R_nrw_*/*R_brd_* as a function of the broadband response. Same data as Figure 5B in the main text. **B** Distribution of the proportional increase *p*(*R_nrw_*/*R_brd_*/*R_nrw_*) in the narrowband response relative to the broadband response, conditioned on different absolute values of the broadband response (colors), as fit by inverse gamma distributions. Arrows in A mark the absolute values of broadband response upon which the proportions are conditioned. Same data as Figure 5C in the main text. **C** Maximum likelihood fits of inverse gamma distributions to the distributions of proportional increase for five absolute values of the broadband response. Small broadband responses tend to be amplified more than large broadband responses. Note that the proportional increase on the x-axis in each of the five panels ranges, respectively, from 1.2x to 33x, 1.2x to 9x, 1.2x to 4.5x, 1.2x to 3.0x, and 1.2x to 2.4x, respectively.

**Figure S6.**
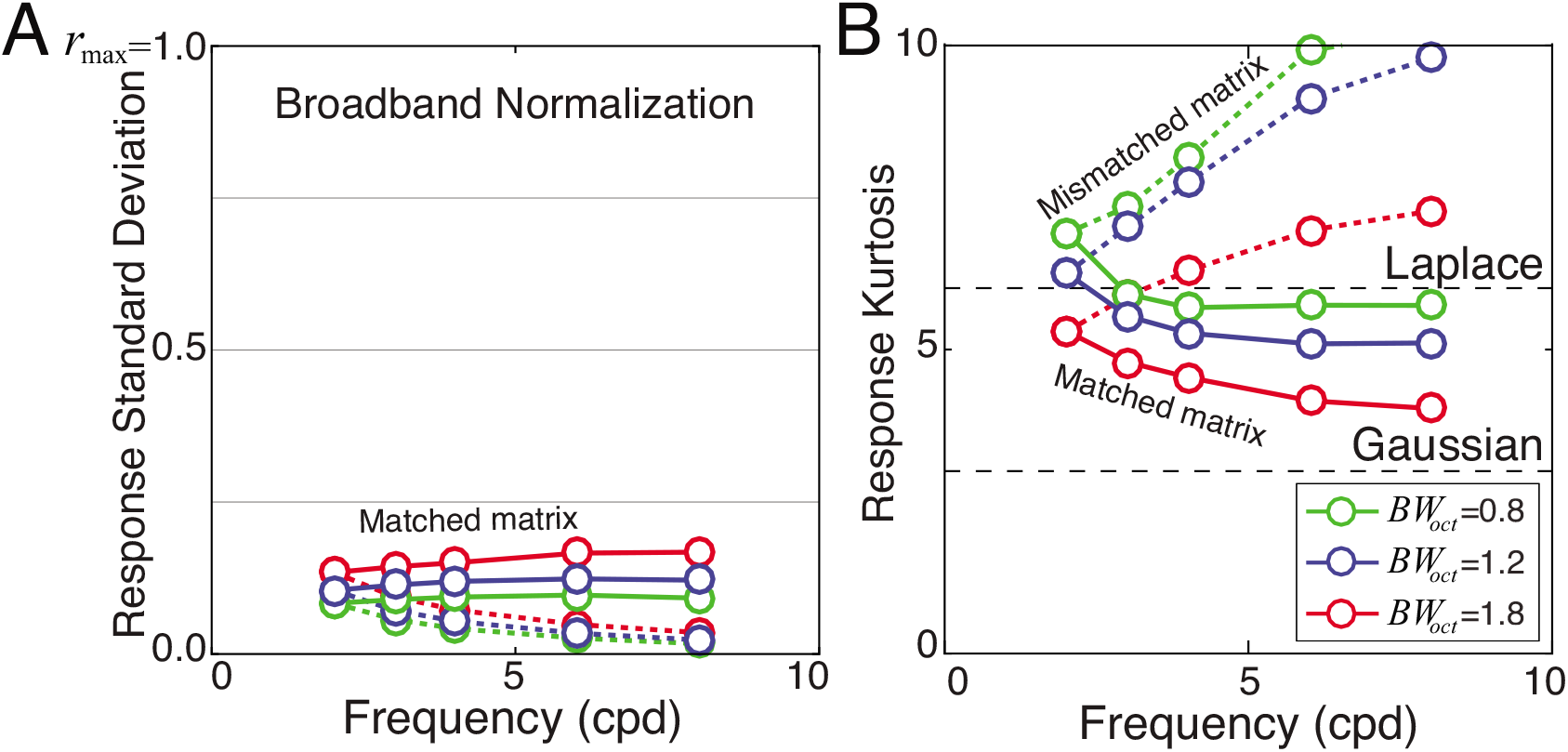
Response statistics with Gabor receptive fields, broadband normalization, and matched and mismatched rectangular weight matrices. **A** Stimulus-driven response standard deviation is invariant to preferred frequency when the receptive field’s weight matrix is matched to the preferred feature (solid curves). Response standard deviation decreases systematically as the magnitude of the mismatch increases (dashed curves). **B** Stimulus-driven response kurtosis is consistent with a Gaussian when the weight matrix is matched to the preferred feature, but increases systematically as spatial frequency (and the magnitude of the mismatch) increases. Decreased stimulus-driven response variation (A) and increased response kurtosis (B) both decrease the signal-to-noise ratio for stimulus discrimination (Fig. S4).

**Figure S7.**
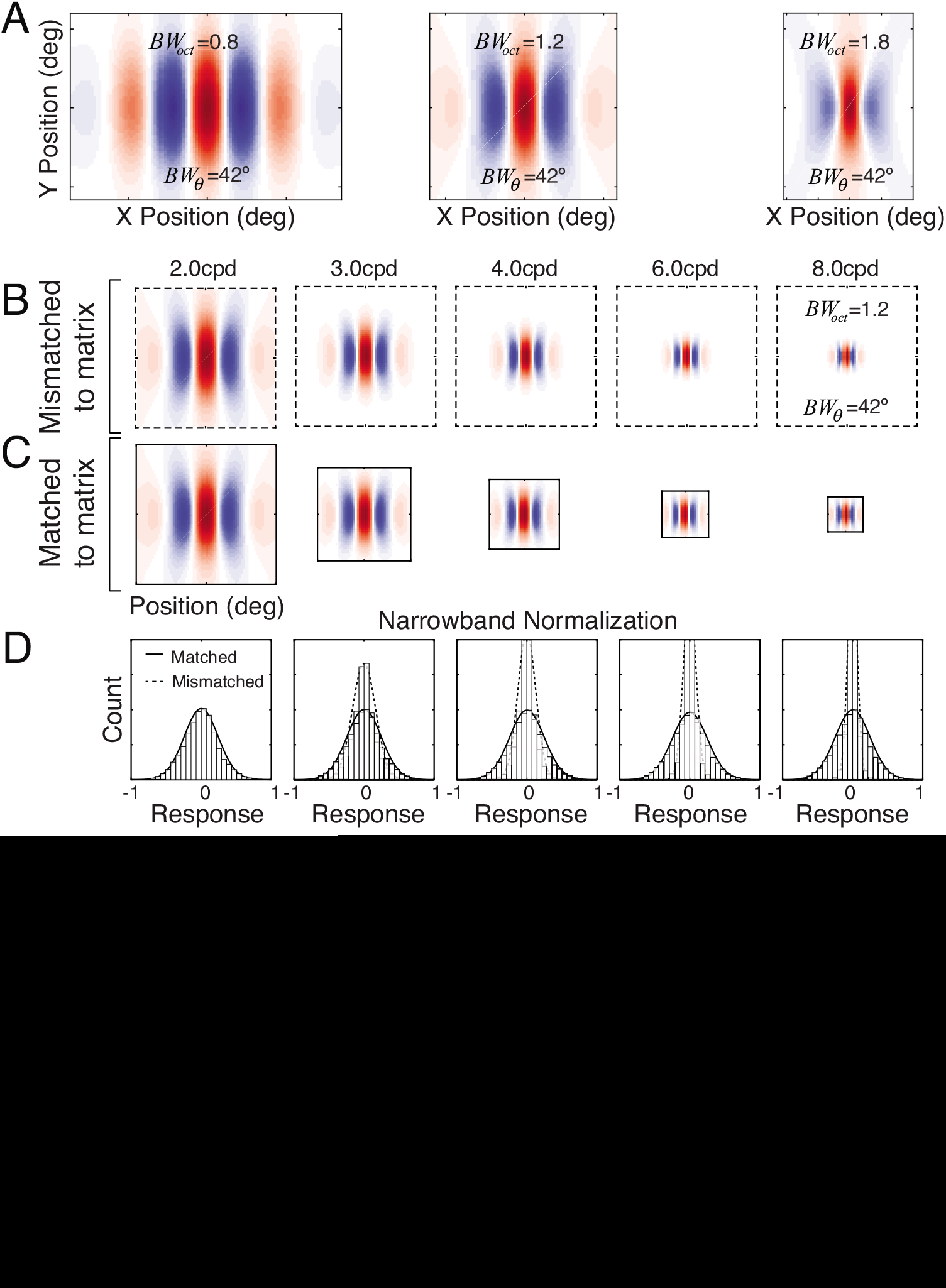
Log-Gabor receptive fields and response statistics. **A** Log-Gabor receptive fields with octave bandwidths of 0.8, 1.2, and 1.8, and orientation bandwidths of 42°. Different octave bandwidths correspond to preferred features with different aspect ratios (see Methods). The receptive fields **B** Mismatched and **C** matched receptive field weight matrices with 1.2 octave bandwidth for five preferred spatial frequencies. **D** Response distributions from matched and mismatched matrices. Matched weight matrices (solid curves) yield response distributions that are invariant to the scale of the preferred feature. Mismatched weight matrices (dashed curves) yield response distributions that change shape and variance with the magnitude of the mismatch. **E** Response standard deviation. Stimulus-driven response variance is constant with preferred frequency when the matrix is matched to the preferred feature. When mismatched, response standard deviation decreases with the magnitude of the mismatch. **F** Response kurtosis is the same as a Gaussian with matched weight matrices, but increases with the magnitude of the mismatch. Log-Gabor and Gabor response statistics are similar (see Fig. 7 in the main text).

**Figure S8.**
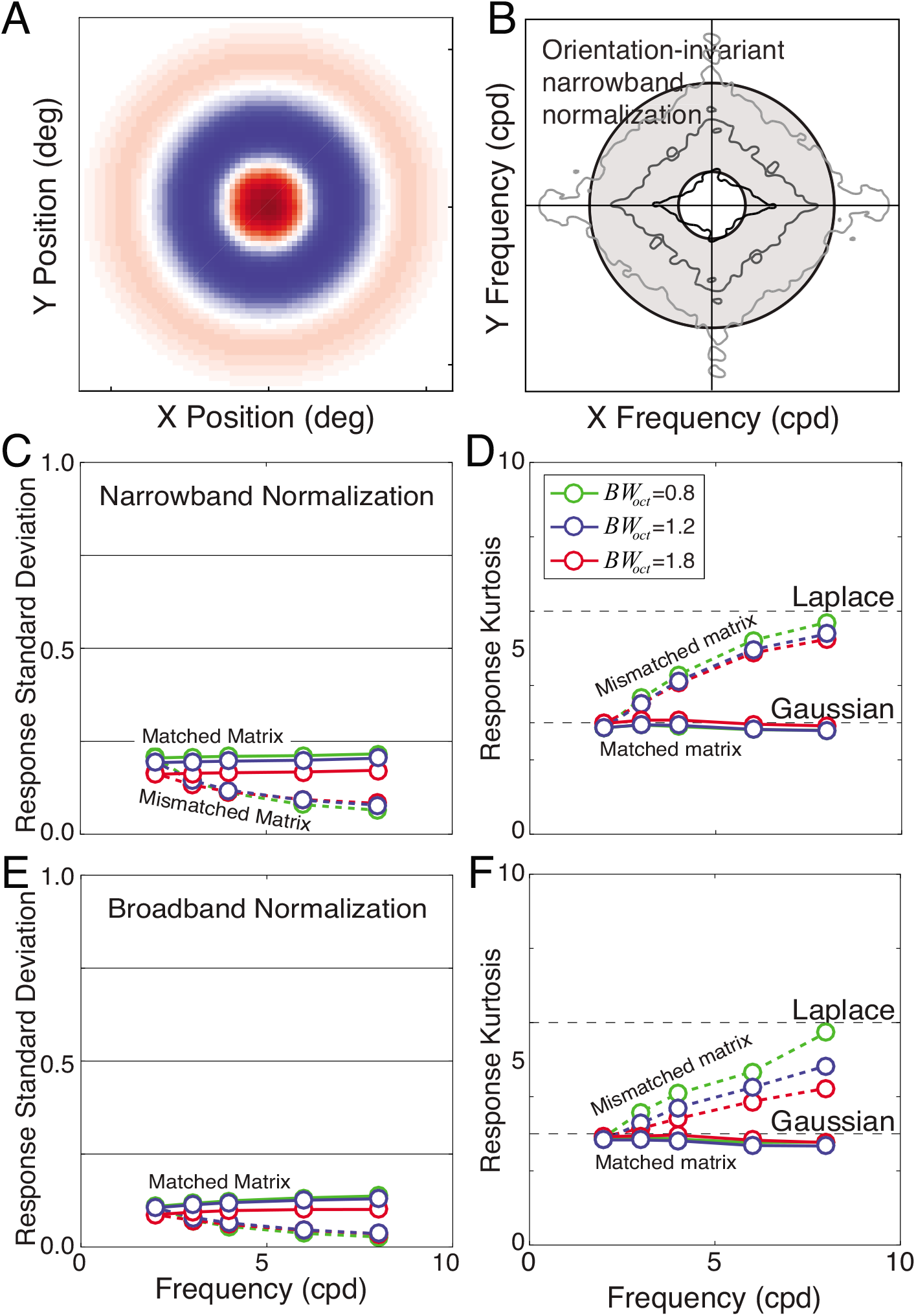
Response statistics for radial Gabor receptive fields with narrowband and broadband normalization. **A** Radially symmetric Gabor receptive field, similar to ganglion cell receptive fields in retina and relay cell receptive fields in lateral geniculate nucleus. **B** Schematic of pooling region in frequency space that determines the narrowband normalization factor. **C** Stimulus-driven response standard deviation with narrowband normalization for three different octave bandwidths (colors). **D** Stimulus-driven response kurtosis with narrowband normalization. **E** Response standard deviation with broadband normalization. **F** Response kurtosis with broadband normalization. Narrowband and broadband response statistics are more similar with radial Gabor receptive fields than with oriented Gabor receptive fields.

**Figure S9.**
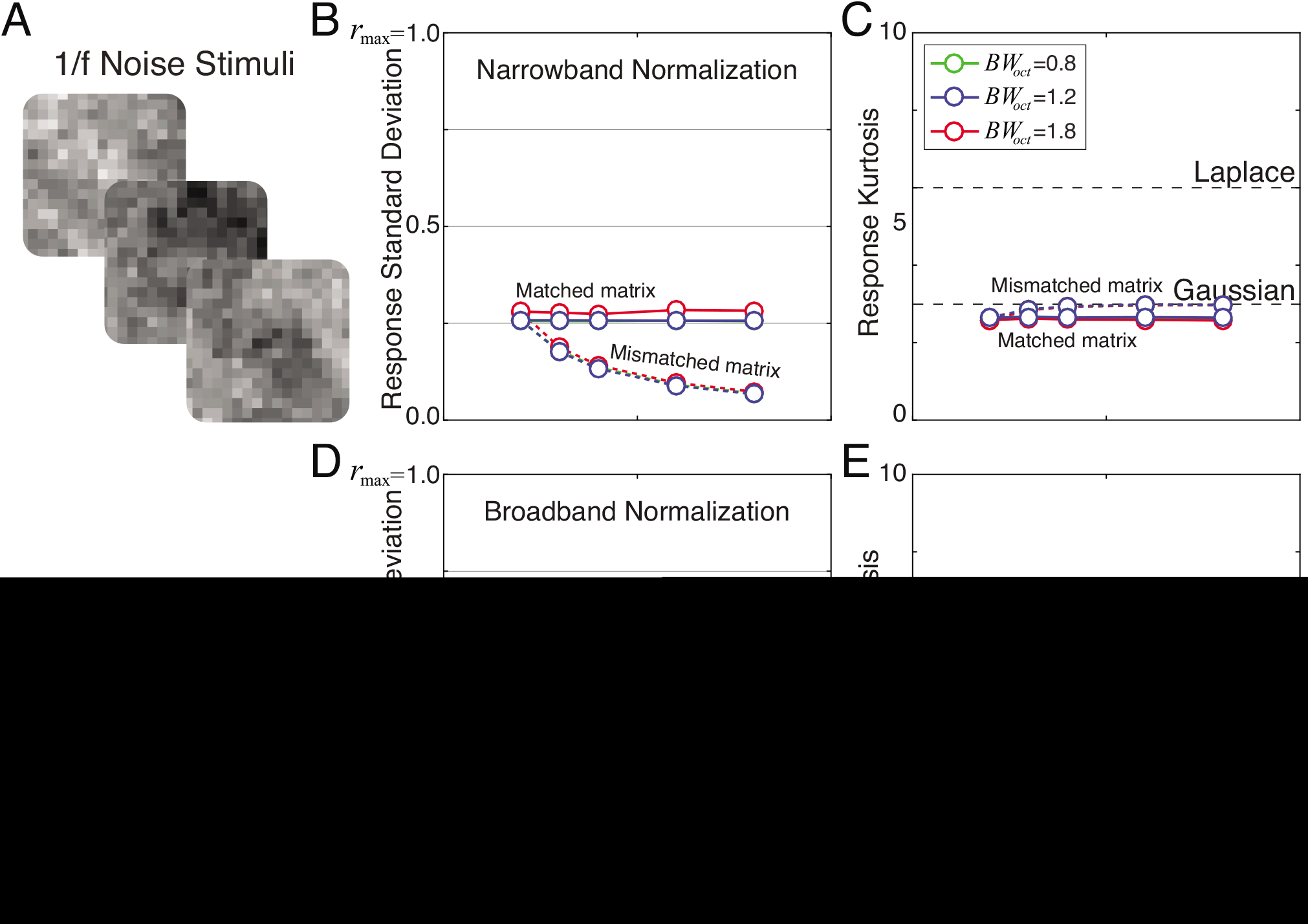
Response statistics to 1/f noise stimuli. Results are plotted for narrowband normalization with matched and mismatched weight matrices. **A** Example 1/f noise stimuli. 1/f noise stimuli were matched in contrast to the natural stimuli. **B** Stimulus-driven response standard deviation with narrowband normalization. **C** Stimulus-driven response kurtosis with narrowband normalization. Regardless of whether the matrices are matched or mismatched, the response kurtosis is consistent with a Gaussian. Note that it may appear surprising that with narrowband normalization, 1/f noise yields similar stimulus-driven response variance as the natural stimuli. (Broadband-normalized responses are substantially smaller with noise than with natural stimuli; see D). This is because our modeling assumes that the normalization factor is computed noiseless. This is not plausible for biological systems. Adding a small constant *N*_0_ to the normalization term changes the response model to *R_nrw_* = **f**^*T*^**c** (*N_nrw_* + *N*_0_) and causes a substantial reduction in the stimulus-driven response variance to noise stimuli while leaving stimulus-driven response variance to natural stimuli relatively unaffected. The small constant has a larger effect on the response variance to noise stimuli because the (noiseless) narrowband normalization factor (i.e. similarity) tends to be much smaller for noise than for natural stimuli. Provided the constant is small enough, its addition to the response model does not appreciably change the distributional shape of the narrowband response distributions. **D** Response standard deviation with broadband normalization. The slight uptick in response standard deviation for the smaller matched matrices vanishes with downsampling. This increase is the only substantive difference we observed with downsampling. **E** Response kurtosis with broadband normalization.

**Figure S10.**
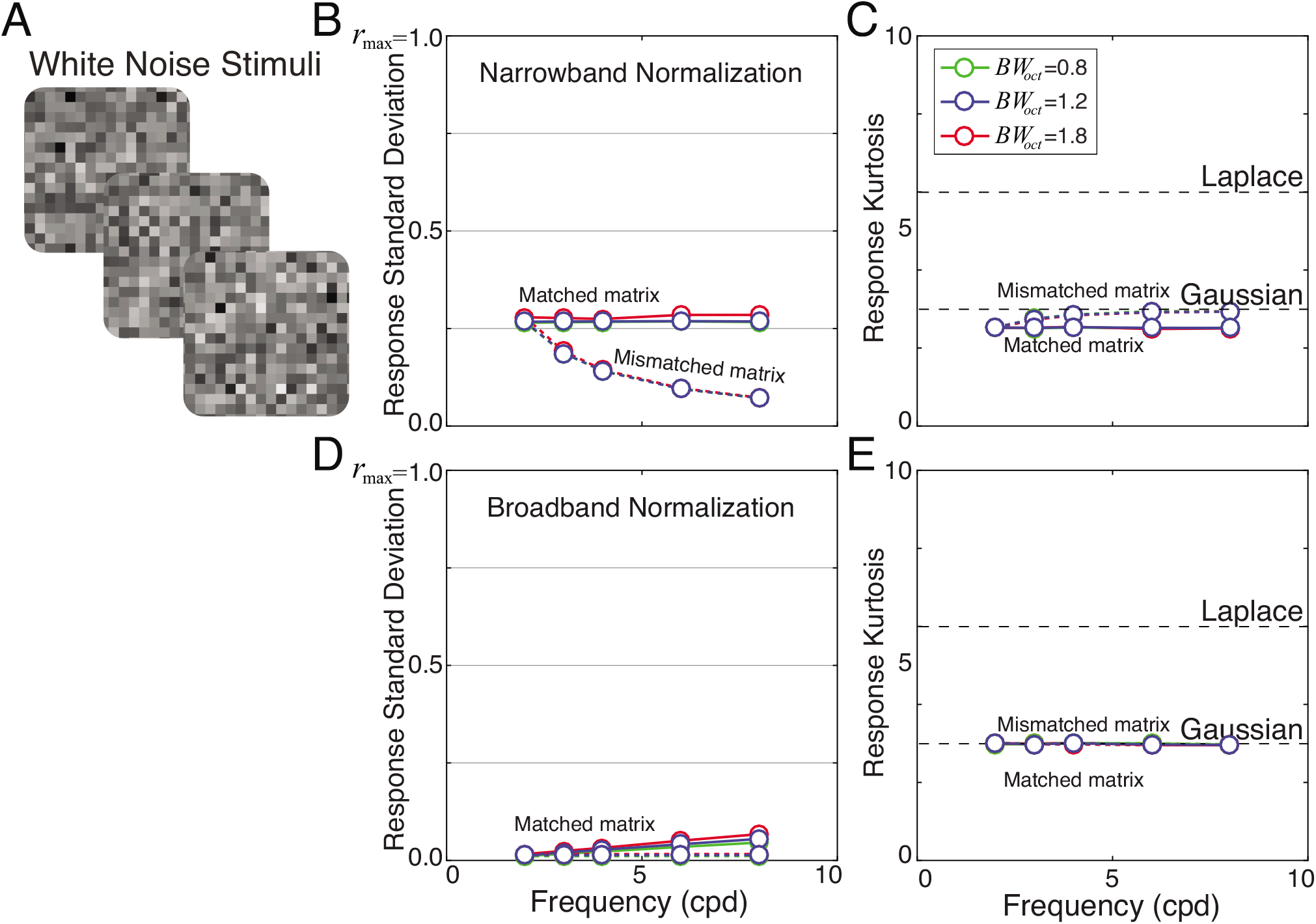
Response statistics to white noise stimuli. Results are plotted for narrowband normalization with matched and mismatched weight matrices. **A** Example 1/f noise stimuli. White noise stimuli were matched in contrast to the natural stimuli. **B** Stimulus-driven response standard deviation with narrowband normalization. **C** Stimulus-driven response kurtosis with narrowband normalization. Regardless of whether the matrices are matched or mismatched, the response kurtosis is consistent with a Gaussian. Note that it may appear surprising that with narrowband normalization, white noise yields similar stimulus-driven response variance as the natural stimuli. (Broadband-normalized responses are substantially smaller with noise than with natural stimuli; see D). This is because our modeling assumes that the normalization factor is computed noiseless. This is not plausible for biological systems. Adding a small constant *N*_0_ to the normalization term changes the response model to *R_nrw_* = **f**^*T*^**c** (*N_nrw_* + *N*_0_) and causes a substantial reduction in the stimulus-driven response variance to noise stimuli while leaving stimulus-driven response variance to natural stimuli relatively unaffected. The small constant has a larger effect on the response variance to noise stimuli because the (noiseless) narrowband normalization factor (i.e. the similarities) tends to be much smaller for noise than for natural stimuli. Furthermore, because the phase-invariant similarity of white noise stimuli is lower than the phase-invariant similarity of 1/f noise stimuli to the receptive field, the constant drives down the stimulus-driven response variance to white noise stimuli more than to 1/f stimuli. Provided the constant is small enough, its addition to the response model does not appreciably change the distributional shape of the narrowband response distributions. **D** Response standard deviation with broadband normalization. The slight uptick in response standard deviation for the smaller matched matrices vanishes with downsampling. This increase is the only substantive difference we observed with downsampling. **E** Response kurtosis with broadband normalization.

**Figure S11.**
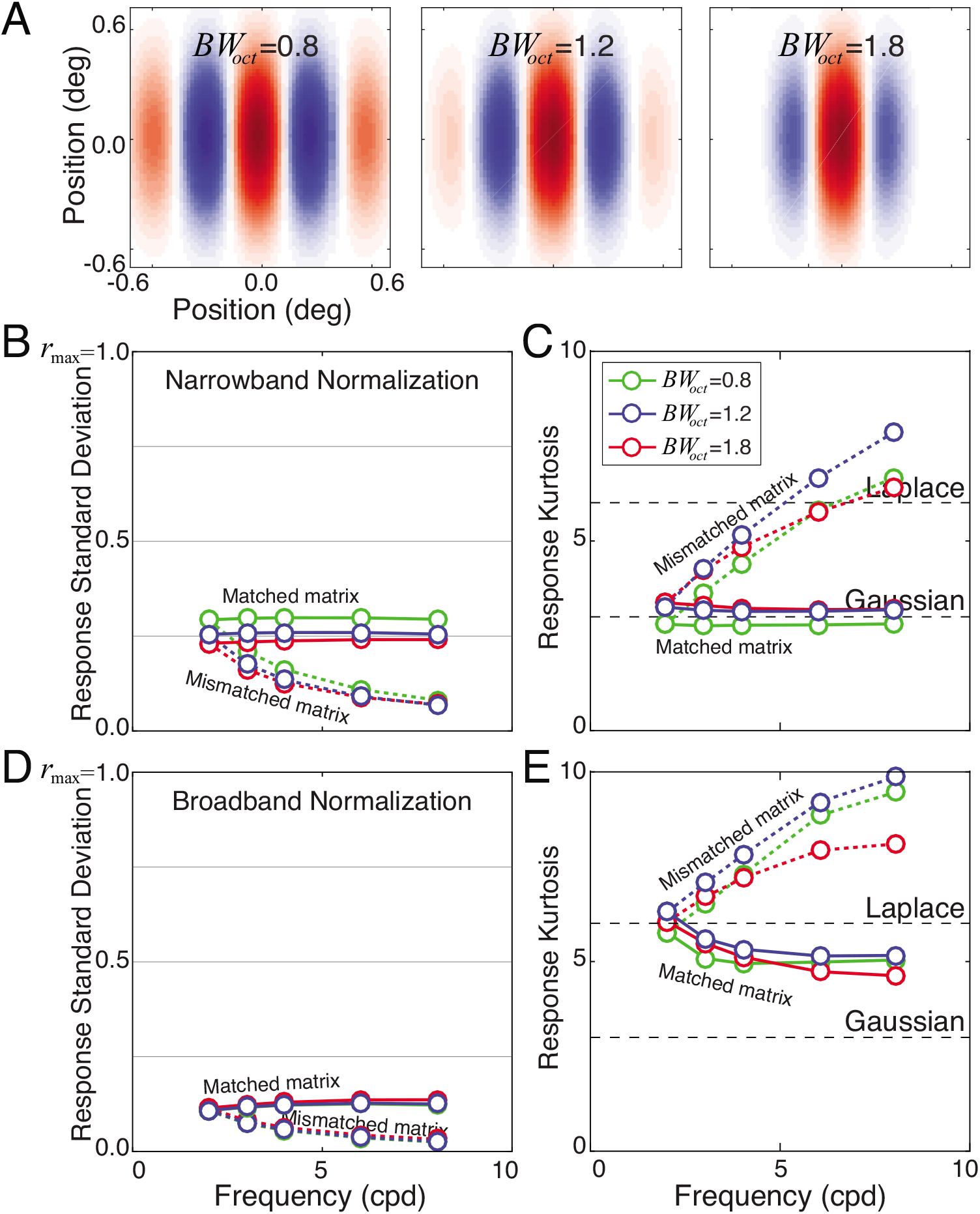
Response statistics with square weight matrices, which are often more convenient for modeling. Preferred features with different octave bandwidths have different aspect ratios. Thus, nominally matched square matrices are actually slightly mismatched for preferred features having octave bandwidths other than 1.2 (see Methods). **A** Preferred features in square weight matrices. **B** Stimulus-driven response standard deviation with narrowband normalization. Results for ‘matched’ square matrices are similar to the results with matched rectangular weight matrices presented in the main text. However, the 0.8 octave bandwidth preferred feature yields slightly higher stimulus-driven response variance and the 1.8 octave bandwidth feature yields slightly lower stimulus-driven variance than their rectangular counterparts. **C** Stimulus-driven response kurtosis with narrowband normalization. Results are very similar with square and rectangular weight matrices. **D** Response standard deviation with broadband normalization. E Response kurtosis with broadband normalization. Square weight matrices yield broadly similar results as rectangular weight matrices, and can probably be used for many applications interchangeably with rectangular matrices.

### Supplement 1 Expected stimulus discriminability for Gaussian response distributions

Consider a neuron whose response can be modeled as a zero-mean Gaussian-distributed random variable *r* with standard deviation *σ_E_* such that

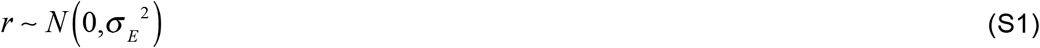

Let *r*_1_ and *r*_2_ be two random response samples. The response difference *u* = *r*_l_ – *r*_2_ is also Gaussian distributed with a variance that is twice the variance of each of the i.i.d. responses

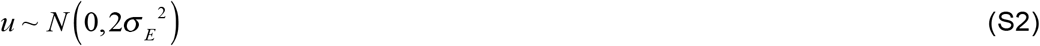

We are interested in the expected absolute difference 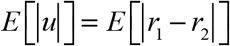 of two random responses. In general, the absolute value of a zero-mean Gaussian distributed random variable σ_2_ obeys a half-normal distribution with mean 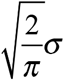.

Given that *u* = *r*_l_ – *r*_2_ is a zero-mean Gaussian variable with variance 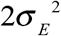, we have

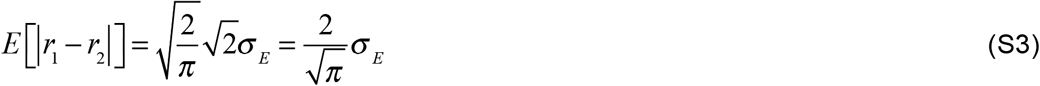

If the neuron’s response is corrupted by encoding noise of variance 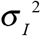, then the expected discriminability across stimuli for this neuron is given by

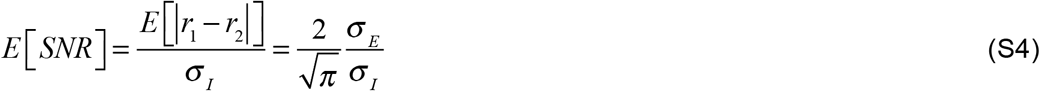

### Supplement 2 Expected stimulus discriminability for Laplace response distributions

Consider a neuron whose response can be modeled as a zero-mean Laplace-distributed random variable *r* with standard deviation *σ_E_* such that

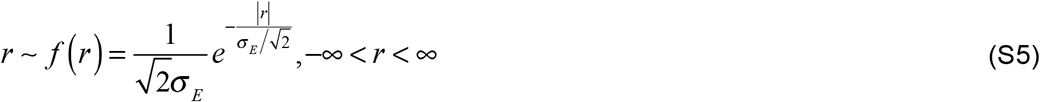

Let *r*_1_ and *r*_2_ be two random response samples. The response difference *u* = *r*_l_ – *r*_2_ is the difference of two i.i.d. responses. Let *u* ~ *g*(*u*)

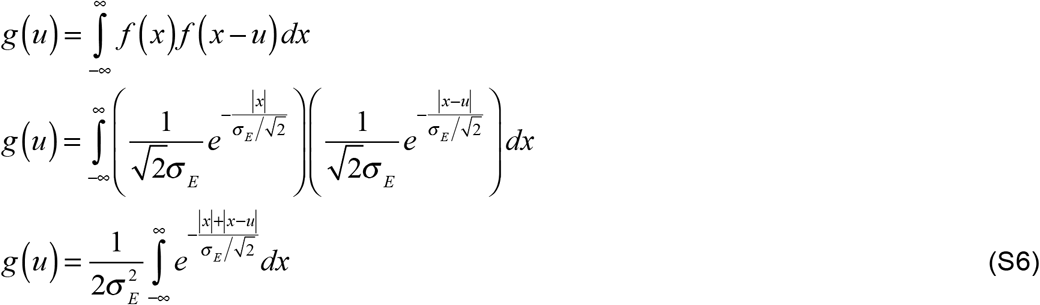

The integral cannot be simply evaluated with an absolute value in the integrand. To remove the absolute value from the integrand, we split the integral depending on the values that *u* and *x* take. We note that *g*(*u*) is even symmetric. Thus, solving the integral for all values of *u* > 0 will provide the solution to the integral for all values of *u <* 0. Assuming that *u >* 0, then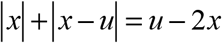 when 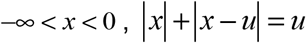 when 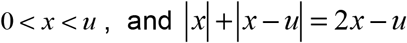 when *u* < *x* < ∞. Evaluating for cases when *u* > 0 yields

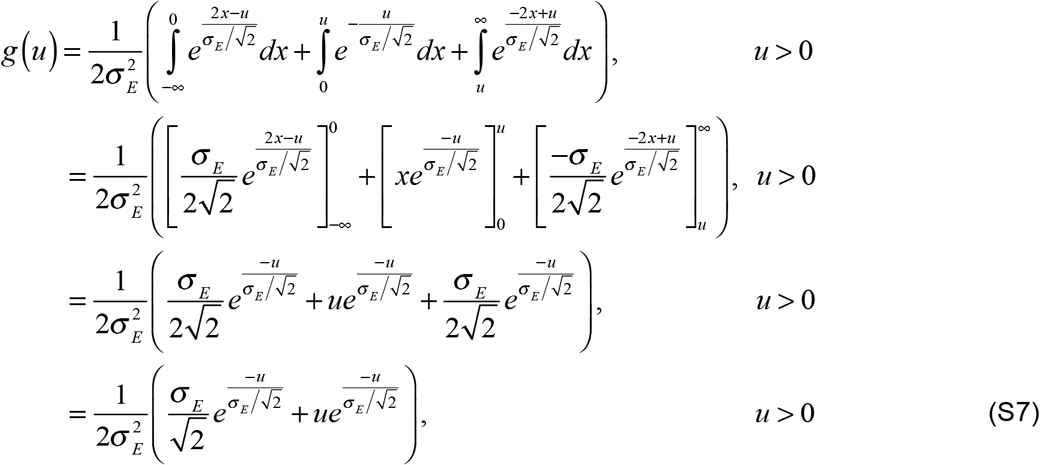

Given that *g*(*u*) is even-symmetric, we can replace *u* with |*u*| in Equation S7

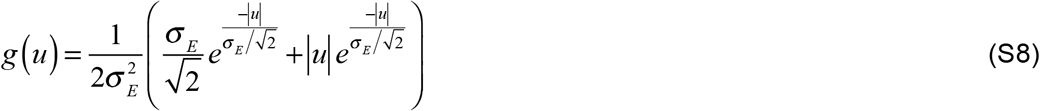

After distributing the leading scale factor 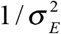 and rearranging terms, we obtain an function where the two terms in the sum are the expressions for a Laplace distribution and a bilateral Gamma distribution

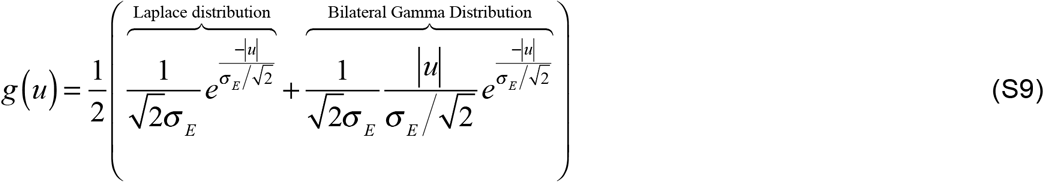

Recall that we are interested in the expectation of the absolute value of |*u*| = |*r*_1_ – *r*_2_|, not the expectation of *u* = *r*_l_ – *r*_2_ itself. Computing the expectation 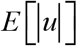 using *g*(*u*) from the definition of expectation

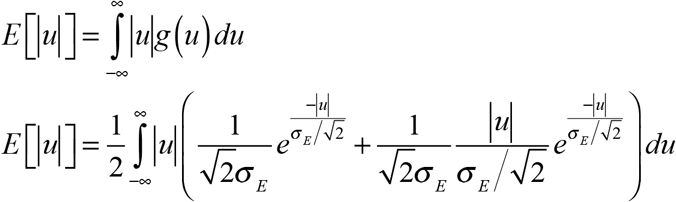

Noting that the integrand is an even function means that twice the integral from zero to infinity equals the integral from negative infinity to infinity

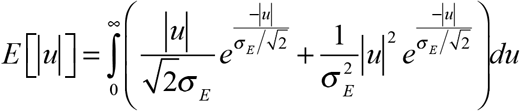

On the positive real line, we can drop the absolute value symbols

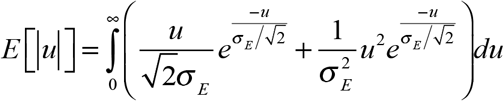

Splitting the integral

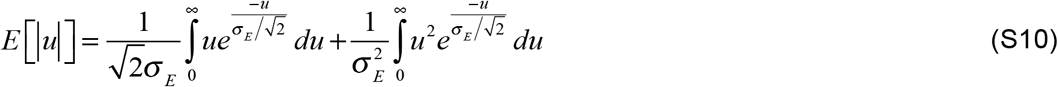

Each of the two definite integrals in equation S6 can be computed with the standard result

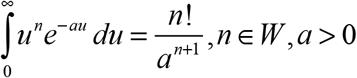

Plugging in

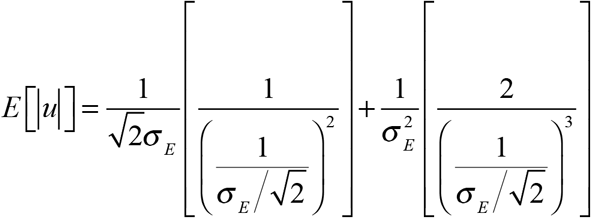

Simplifying terms

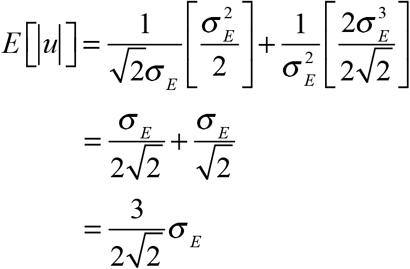

Therefore, the mean absolute difference between two i.i.d. mean-zero Laplace random variables of standard deviation *σ_E_* is

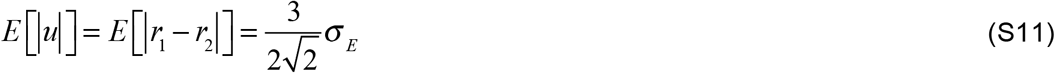

For internal noise of standard deviation *σ*_1_, the expected stimulus discriminability across all stimuli is given by

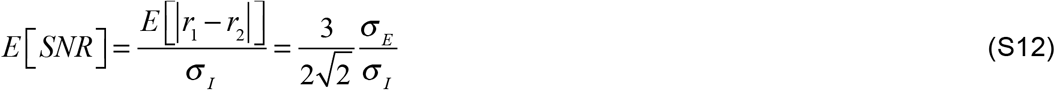

